# BRD9-SMAD2/3 orchestrates stemness and tumorigenesis in pancreatic ductal adenocarcinoma

**DOI:** 10.1101/2023.03.02.530770

**Authors:** Yuliang Feng, Liuyang Cai, Martin Pook, Feng Liu, Chao-Hui Chang, Mai Abdel Mouti, Reshma Nibhani, Shihong Wu, Siwei Deng, Stefania Militi, James Dunford, Martin Philpott, Yanbo Fan, Guo-Chang Fan, Qi Liu, Jun Qi, Sakthivel Sadayappan, Anil G. Jegga, Udo Oppermann, Yigang Wang, Wei Huang, Lei Jiang, Siim Pauklin

## Abstract

The dismal prognosis of pancreatic ductal adenocarcinoma (PDAC) is linked to the presence of pancreatic cancer stem-like cells (CSCs) that respond poorly to current chemotherapy regimens. By small molecule compound screening targeting 142 epigenetic enzymes, we identified that bromodomain-containing protein BRD9, a component of the BAF histone remodelling complex, is a key chromatin regulator to orchestrate the stemness of pancreatic CSCs via cooperating with the TGFβ/Activin-SMAD2/3 signalling pathway. Inhibition and genetic ablation of BDR9 block the self-renewal, cell cycle entry into G0 phase and invasiveness of CSCs, and improve the sensitivity of CSCs to gemcitabine treatment. In addition, pharmacological inhibition of BRD9 significantly reduced the tumorigenesis in patient-derived xenografts mouse models and eliminated CSCs in tumours from pancreatic cancer patients. Mechanistically, inhibition of BRD9 disrupts enhancer-promoter looping and transcription of stemness genes in CSCs. Collectively, the data suggest BRD9 as a novel therapeutic target for PDAC treatment via modulation of CSC stemness.

## Introduction

Pancreatic cancer with its most common type, pancreatic ductal adenocarcinoma (PDAC), is one of the most lethal human malignancies ^1-3^. It has an overall median survival time of 6-9 months and a similarly 5-year survival rate of 6%, making it currently the fourth leading cause of cancer-related deaths in western countries^4 5^. Due to the increasing incidence of risk factors including obesity and other metabolic traits, pancreatic cancer is projected to overtake colorectal, breast and prostate cancer and become the second leading cause of worldwide cancer-related deaths by 2030^6^. The disease owes its exceptional level of lethality to multiple factors. Pancreatic cancer often has no early symptoms and presents itself in an advanced stage at diagnosis: only 20% of newly diagnosed pancreatic cancers are amenable to surgery^7^. In turn, the disease’s poor response to chemotherapy and radiotherapy results in disease re-emergence for 90% of the surgically treated patients^8^. Treatment options for PDAC are limited and inefficient. Gemcitabine, an antimetabolite drug of the nucleoside analogue class is the standard of care in PDAC therapy^8^. It is currently the only approved single-agent drug for pancreatic cancer, with still modest improvements in survival rate^9^, whether the drug is administered alone or in combination with adjuvant drugs such as the EGFR inhibitor erlotinib^10^ or the tubulin-targeting drug Nab-paclitaxel^9^. A combination therapy such as FOLFIRINOX is superior to gemcitabine-based regimens in restraining the progression of metastatic PDAC but has lower tolerability^11^.

Precancerous lesions and dedifferentiation of the cells to a progenitor-like or stem cell-like state with increased cellular plasticity frequently occur during pancreatic tissue transformation^4 12^. A distinct cell population often referred to as cancer stem cells (CSCs), seems to acquire a stem cell-like state partially resembling naturally occurring stem cells^13-15^. This phenotype allows them to give rise to the whole tumour with its entire cellular heterogeneity and thereby supports metastases formation and development of resistance to current cancer therapeutics. The existence of developmentally plastic cancer stem cells has been discovered in the brain, breast, colon, oesophagus, liver, lung, ovarian, prostate, stomach and thyroid cancers, among others. In the case of PDAC, the first reports of cancer stem cells date back to 2007^13 14^. Since then, pancreatic CSCs have been conclusively shown to be involved in PDAC resistance to chemotherapy, displaying increased prevalence within the tumour after treatment with gemcitabine^16 17^. Annihilating CSCs is thus emerging as an essential aim of PDAC therapeutics. CSCs are thought to have specific epigenetic mechanisms^18^ that regulate their self-renewal, and the formation of CSCs has been postulated to occur as a result of epigenetic events^19^. Accordingly, cancer epigenetics has established itself as a promising area of oncology research^20^. After a decade in which epigenetic cancer drugs were approved only for haematological malignancies, the first FDA approval for an epigenetic drug targeting solid tumours was granted in 2020 for an EZH2 inhibitor. Despite their paradigm-shifting novel mechanism of action, early results of epigenetic modulators have identified the need for better selection of targets, improved intratumoral drug penetration and elimination of CSCs.

Pancreatic cancers are complex tumours with significant heterogeneity in their molecular and cellular make-up that are controlled by various signalling pathways that crosstalk with epigenetic regulators. Among these pathways is the TGF β/Activin/Nodal-SMAD2/3 pathway. This developmental signalling pathway plays a central role in early development by regulating the self-renewal of human pluripotent stem cells (hPSCs), Epithelial-to-mesenchymal transition (EMT) and pancreatic tissue homeostasis^21-23^. The TGFß/Activin/Nodal-SMAD2/3 pathway regulates epigenetic mechanisms, for instance by cooperating with the core pluripotency protein NANOG and epigenetic modifiers such as DPY30-COMPASS to control pluripotency and differentiation of hPSCs^24^. Aside from the key role of TGF β/Activin/Nodal-SMAD2/3 in pluripotent stem cells and developmental processes, this signalling pathway is directly involved in the formation of PDAC^22^ and is frequently deregulated in PDAC^25 26^. The function of the pathway in PDAC is particularly interesting, because it confers dedifferentiated stem cell-like features to CSCs in PDAC^15^, although the underlying mechanisms are still largely unknown.

Our hypothesis was that epigenetic or chromatin-templated mechanisms are essential to main CSC properties and that inhibition of these mechanisms may provide a path forward to modulate CSC phenotypes. Using a focused compound library of epigenetic inhibitors we performed a small molecule compound screening and identified the BAF chromatin complex component BRD9 as a critical regulator of CSC behaviour. Our results indicate that BRD9 is an attractive therapeutic target for specifically eliminating CSCs in PDACs.

## Results

### Development of a screening platform to target PDAC CSCs

Since PDAC cells comprise a heterogeneous population of CSCs and non-CSCs, we first decided to establish a suitable small molecule screening platform in PDAC cells (**Figure 1A**). To uncover novel regulators of CSCs, we used three CSC markers (OCT4, CD133, SSEA4) for identifying stem cell-like cells in the experiments. OCT4 is a transcription factor crucial to the self-renewal and pluripotency of embryonic stem cells^27^ and is associated with worse outcomes in PDAC^28^. Besides OCT4, we used two other CSC markers: surface protein CD133 and glycolipid carbohydrate epitope SSEA4. A high expression of CD133 in PDAC patients (over 5%) has been linked to a 0% 5-year patient survival rate, compared to 23.5% for CD133-negative tumors^29^, and PDAC metastasis was found to depend upon a subset of CD133-expressing cells^13^ which trigger early metastasis to lymph nodes^29^. Finally, the third stem cell marker, stage-specific embryonic antigen-4 (SSEA4), is known to be expressed in a wide range of cancers, including PDAC^30^. The pancreatic cancer cells used for this screening were genetically engineered, with the sequence coding for green fluorescent protein (eGFP) integrated in the endogenous locus via TALEN-mediated recombination, resulting in controlled expression of a OCT4-eGFP fusion protein driven by the endogenous *OCT4* promoter (**Supplemental Figure 1A**). To validate the importance of CD133, SSEA4 and OCT4 as markers for the CSC population in our PDAC cells, we treated the cells with chemotherapy reagents gemcitabine (GEM), paxclitaxel (PAX) and 5-fluorouracil (5-FU) that are currently in clinical use as PDAC patient therapeutics. Gemcitabine, paxclitaxel and 5-fluorouracil treatment of PDAC cells for 5 days enriched for cells expressing OCT4-GFP, CD133 and SSEA4 from ∼1% triple positive cells to ∼75%, ∼25% and ∼60%, respectively (**Supplemental Figure 1B**), by eliminating most of the PDAC cells that do not express these three CSC markers (**Supplemental Figure 1C**). This results in selective survival and enrichment of the rare OCT4-GFP+/CD133+/SSEA4+/EPCAM+ CSCs.

**Figure 1.**
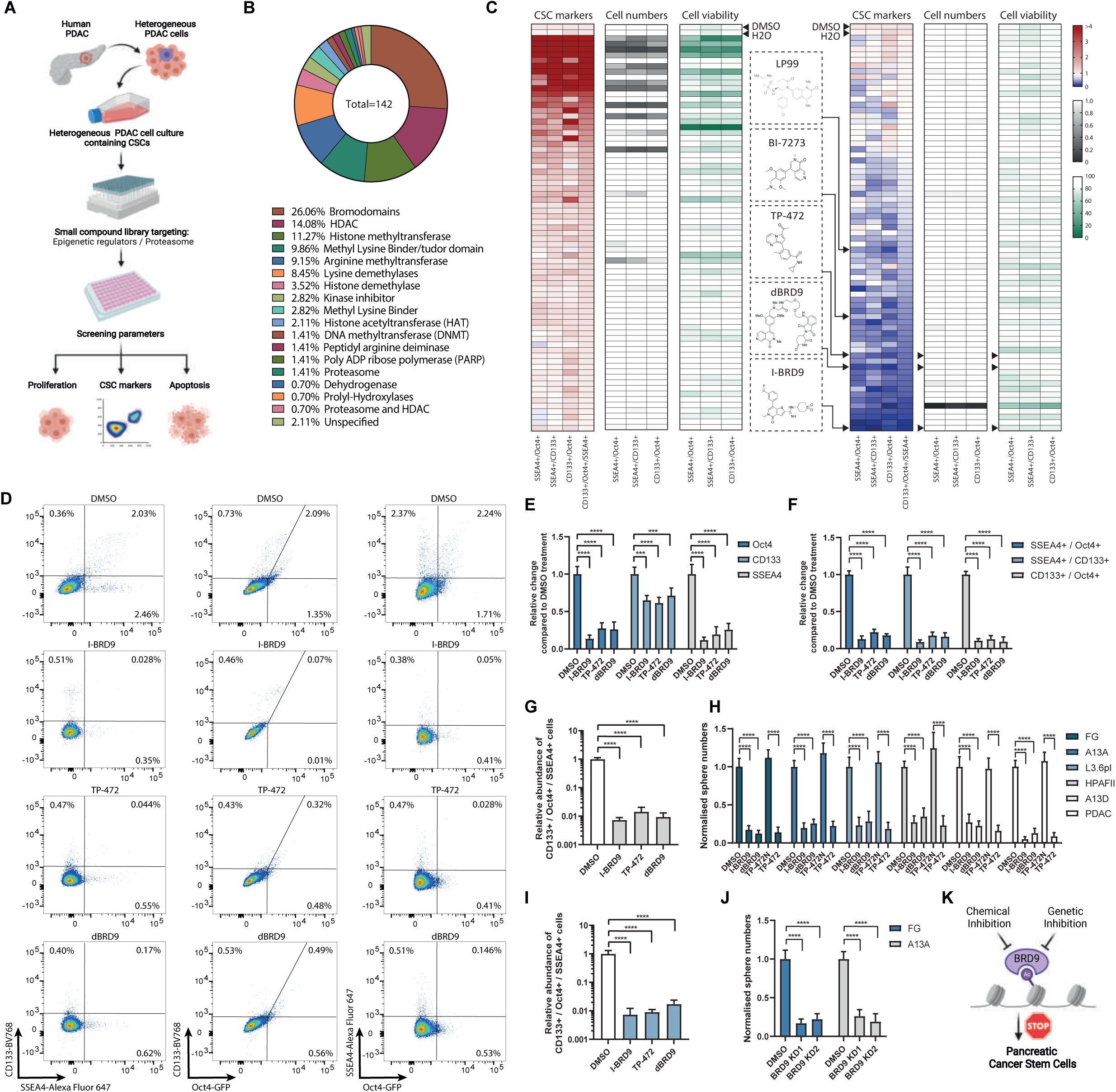
Chemical screening identifies BRD9 as a regulator of pancreatic CSCs. (**A**) Schematic depiction of the small molecule compound screening process on pancreatic cancer cells. (**B**) The classification of the compound library based on the percentage of the 142 compounds belonging to each class of enzymes they target. (**C**) BRD9 inhibitors decrease the relative number of cells that express CSC markers OCT4-GFP, CD133 and SSEA4 as double positive cells or triple positive cells. Heat maps of chemical screening depicting the relative change in the expression of CSC markers, cell numbers and cell viability. Various BRD9 inhibitors, I-BRD9, dBRD9, TP-472, BI-7273, LP99, are marked in the heat map results. (**D**) BRD9 inhibitors reduce the percentage of OCT4-GFP, CD133 and SSEA4 double positive cells. Representative dot blot graphs from flow cytometry analyses of CSC markers in pancreatic cancer cells. (**E-G**) BRD9 chemical inhibitors reduce OCT4-GFP, CD133 and SSEA4 marker expression in pancreatic cancer cells. The relative change in single CSC marker positive (**E**), double positive (**F**), and triple positive cells compared to control DMSO treatments (**G**). (**H**) BRD9 inhibition reduces pancreatic CSC self-renewal in different PDAC cell lines and PDAC cells from surgically resected tumour. The impact of BRD9 inhibition during tumour sphere formation as quantified by the changes in the number of spheres. (**I**) The relative decrease in OCT4-GFP+/CD133+/SSEA4+ triple positive cancer cells by BRD9 inhibition in L3.6pl PDAC cell line. (**J**) BRD9 knockdown reduces pancreatic CSC self-renewal in PDAC cell lines. BRD9 stable knockdown was performed by BRD9-specific shRNAs compared to Scramble control. (**K**) Schematic depiction of the effects of BRD9 inhibition in pancreatic CSCs. The chemical inhibition and genetic inhibition of BRD9 reduces pancreatic CSCs. Experiments represent three replicates. Statistical analysis was performed by 2-way ANOVA with multiple comparisons with Tukey correction and **** marks adjusted P-value <0.0001, *** is adjusted P-value <0.001, ** is adjusted P-value <0.01, * is adjusted P-value <0.05.

Since the TGFβ Activin/Nodal-SMAD2/3 developmental signalling pathway regulates pluripotency via epigenetic regulatory complexes and impacts stem cell-like characteristics of CSCs^24 31^, we also tested the impact of TGF β/Activin signalling on CSC resistance to currently used chemotherapeutics by treating the cells with the TGF β/Activin signalling inhibitor SB431542 in combination with Gemcitabine, Paxclitaxel and 5-FU (**Supplemental Figure 1B-C**). Inhibition of TGF β/Activin signalling strikingly reduced the chemoresistance of PDAC cells as indicated by reduced numbers of OCT4-GFP+/CD133+/SSEA4+ CSCs (**Supplemental Figure 1B**) and therefore the overall number of surviving PDAC cells (**Supplemental Figure 1C**). These results emphasize the crucial importance of TGF β/Activin signalling on CSC maintenance and their elevated chemoresistant characteristics. Furthermore, PDAC patients with high expression of CD133 and OCT4 have lower overall survival (**Supplemental Figure 1D**) and lower disease-free survival (**Supplemental Figure 1E**). These results underline the suitability of our screening platform and the translational relevancy of this preclinical model for compound screening.

### A focused compound library screen of epigenetic inhibitors identifies BRD9 as an important factor for governing CSC characteristics

Due to the importance of epigenetic pathways in tumorigenesis, we hypothesized that the formation and maintenance of pancreatic CSCs are controlled by epigenetic mechanisms, and these could be used as therapeutic targets for eliminating CSCs. To address this, we performed a focused library screen consisting of validated small molecule inhibitors (**Supplementary Table 1**) targeting epigenetic regulators such as ‘readers, writers and erasers’ of a histone code^32^. These experiments aimed to identify molecular targets of small molecule compounds which specifically affect CSC marker-expressing cells (**Figure 1A-C**). We measured CSC marker expression, cell growth and cell death by flow cytometry (CD133+/OCT4+/SSEA4+) after incubating the cells with the compounds for 5 days, which allowed us to identify effective compounds that impact CSC marker expressing populations while also detecting cells that do not express these CSC markers. The compound library used in our experiments consisted of 142 compounds that have been verified to be active and targeting specific epigenetic modifying enzymes (**Figure 1C**). A substantial fraction contained in the library were acetyl lysine (bromodomain) inhibitors that represented approximately 26% of the compounds, followed by histone deacetylase (HDAC) inhibitors (14%), Histone methyltransferase inhibitors (11.3%), compounds targeting methyl lysine binders/tudor domains (9.9%), arginine methyltransferases (9.2%) and lysine demethylases (8.5%). Overall, this screening identified compounds that significantly and reliably reduced the relative percentage of triple+ marker (CD133+/OCT4+/SSEA4+) expressing CSCs and also reduced cancer cell survival (**Figure 1C**). Our study identified the BET bromodomain inhibitors I-BET 672 and JQ1 as candidate compounds, both of which have previously shown antitumour activity^33^ and synergy with gemcitabine^34^, confirming the suitability of our screening platform.

Importantly, the screening identified novel compounds that target distinct epigenetic regulatory components and strikingly reduced the percentage of CSCs. Among the top candidate compounds with a distinct effect on CSCs we identified BRD9 inhibitors (I-BRD9, dBRD9, TP-472, BI-7273, LP99) while the corresponding inactive negative control compound (TP-472N) did not reduce CSC marker expression (**Figure 1C-D**). These results suggested that BRD9 inhibition could induce differentiation of therapy-resistant CSCs into a more therapy-responsive population and thus could possibly sensitize PDAC cancers to conventional therapy. BRD9 is a component of the non-canonical BAF chromatin remodelling complex (ncBAF)^35^ and has recently been shown to constitute a barrier in reprogramming of somatic cells to induced pluripotent stem cells^36^. The catalytic component SMARCA4 of the BAF complex regulates stem cell properties in pediatric gliomas and provides a possible therapeutic angle in glioblastoma treatment^37^. Collectively these data provided evidence to focus our investigation on the molecular role of the BAF complex and BRD9 in particular, in PDAC CSCs.

To further validate BRD9 as a target we used two BRD9 chemical inhibitors (I-BRD9, TP-472) and the PROTAC degrader of BRD9 (dBRD9)^38^ to investigate their effects on CSC markers (**Supplemental Figure 1F-G**). All three inhibitors of BRD9 resulted in the reduction of OCT4, CD133 or SSEA4 single marker positive cells (**Figure 1E**), even more pronounced reduction in double positive cells for OCT4+/SSEA4+, CD133+/SSEA4+ or CD133+/OCT4+ (**Figure 1F**), and with a most striking reduction in OCT4+/CD133+/SSEA4+ triple positive PDAC cells (**Figure 1G**). I-BRD9 and TP-472 reduced the OCT4+/CD133+/SSEA4+ triple positive CSC numbers also in other PDAC cell lines (**Figure 1I**).

The bulk cell population of the PDAC is sensitive to chemotherapy while the CSC subpopulation of PDAC cells is the reason why PDAC is therapy recalcitrant. Therefore, we investigated the effects of compounds on CSCs by 3D tumour sphere assays. To study the impact of BRD9 inhibition on CSC self-renewal capacity we performed tumour sphere assays with three different BRD9 inhibitors (I-BRD9, TP-472, dBRD9) in five different PDAC cell lines and primary cells from surgically resected PDAC tumours (**Figure 1H**). Inhibition of BRD9 with these three inhibitors showed a significant reduction in CSC sphere numbers while the negative control compound of TP-472 (TP-472N) did not have such effects in any of the cell lines or tumour-derived cells. To investigate the effect of BRD9 inhibition by genetic means, we performed stable BRD9 knockdown (KD) in two PDAC cell lines (**Supplemental Figure 1H-K**). Tumour sphere assay revealed a reduction of spheres in both cell lines by two different shRNA constructs (**Figure 1J**), indicating that BRD9 loss-of-function via knockdown phenocopies the reduction of CSC self-renewal as seen with chemical inhibition of BRD9 (**Figure 1K**).

### BRD9 inhibition reduces pancreatic CSCs entry to G0 cell cycle phase and chemoresistance

Since CSCs have lower sensitivity to chemotherapeutics besides their self-renewing capacity, we investigated the impact of BRD9 inhibition on the chemoresistance of CSCs in combination with Gemcitabine treatment (**Figure 2A**). BRD9 inhibition together with Gemcitabine treatment led to a stronger reduction in CSC sphere formation compared to either I-BRD9 or Gemcitabine treatment alone, suggesting that BRD9 inhibition sensitizes CSCs for Gemcitabine-mediated cell killing. The TGFβ/Activin pathway inhibitor SB431542 reduced the number of CSCs compared to Activin A treated cells (**Supplemental Figure 1L-M**), indicating that the TGFβ/Activin pathway promotes CSC characteristics. To understand whether BRD9 inhibition engages in crosstalk with TGFβ/Activin signalling, we co-treated the cells with I-BRD9 and Gemcitabine along with Activin A or the TGFβ/Activin pathway inhibitor SB431542 during tumour sphere formation. BRD9 inhibition as well as TGFβ/Activin pathway inhibition strikingly increased the sensitivity of CSCs to Gemcitabine (**Figure 2B-C**), while the combination of I-BRD9 and Gemcitabine treatment together with TGFβ/Activin pathway inhibition very efficiently eliminated the formation of spheres (**Figure 2B, D**; **Supplemental Figure 1N**). The effect of BRD9 inhibition on sensitizing cells to Gemcitabine was further validated by three different BRD9 inhibitors (**Supplemental Figure 1O**). Furthermore, BRD9 genetic knockdown in two different PDAC lines reduced CSC chemoresistance not only for Gemcitabine but also 5-FU (**Supplemental Figure 1P-R**).

**Figure 2.**
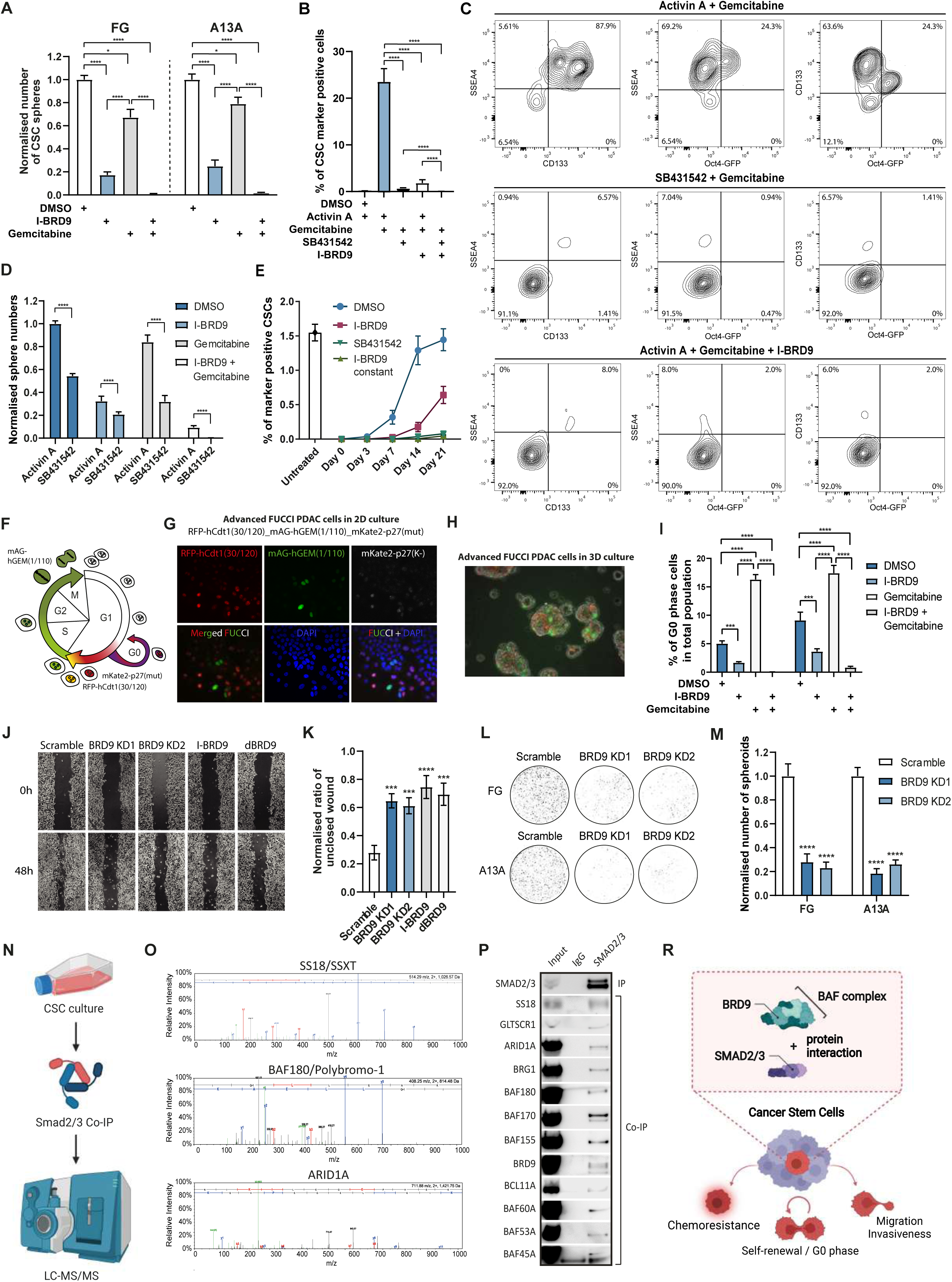
BRD9 inhibition abolishes CSCs characteristics and blocks cells from entering the G0 phase. (**A**) BRD9 inhibition sensitizes CSCs for Gemcitabine mediated elimination. Tumour sphere assays on cells treated with I-BRD9, Gemcitabine or by both simultaneously. (**B-C**) Triple positive OCT4-GFP+/CD133+/SSEA4+ CSC marker reduction in FG cells is induced by Gemcitabine co-treatment with I-BRD9 or SB431542 as shown in bar graph (**B**) or density maps from flow cytometry analyses (**C**). (**D**) TGFβ ctivin signalling inhibition and BRD9 inhibition impair CSC self-renewal and sensitize CSCs for Gemcitabine-mediated destruction. (**E**) BRD9 inhibition shift the balance between CSCs and non-CSCs toward non-CSCs. The cells were analysed for the percentage of OCT4-GFP+/CD133+/SSEA4+ triple positive cells. For I-BRD9 pre-treated sample the cells were treated for 5 days with I-BRD9 before cells sorting, for the I-BRD9 constant and SB431542 constant samples, the cells were treated for 5 days either I-BRD9 or SB431542, the cells were then sorted to OCT4-GFP-/CD133-/SSEA4-negative cells and grown in their constant presence for the rest of the experiment. Untreated cells represent cells that were not treated and not sorted. (**F**) Schematic depiction of the three-colour cell cycle analysis system which enables distinguishing between the cells in early G1, late G1, early S, S/G2/M and G0 phases. (**G**) Fluorescence microscopy of 3FUCCI-PDAC cells grown in 2D condition. (**H**) Fluorescence microscopy overlayed with bright field image of 3FUCCI-PDAC cells grown in 3D sphere condition. (**I**) BRD9 inhibition blocks cells from entering the G0 phase. 3FUCCI-PDAC spheres were treated with I-BRD9, Gemcitabine or co-treated for 3 days followed by flow cytometry analysis. (**J-K**) BRD9 inhibition reduces cell migration. Bright field images of wound healing assay at 0h and 48h time points (**J**) normalised data of wound healing assay as a bar graph. (**L-M**) BRD9 inhibition reduced PDAC cell invasiveness. BRD9 knockdown cells or Scramble cells were used for cell invasiveness assays (**L**) and the results were quantified as normalised number of cells that have undergone matrix invasion (**M**). (**N**) Schematic overview of the SMAD2/3 proteomic experiment. CSCs stimulated with Activin A for 24h and the nuclear fraction of CSCs were used SMAD2/3 co-immunoprecipitation experiments followed by mass-spectrometry analyses. (**O**) SS18/SSXT, BAF180/Polybromo-1 and ARID1A form a complex with SMAD2/3 in CSCs. Peptides corresponding to SS18, BAF180 and ARID1A were identified by mass-spectrometry analyses. (**P**) SMAD2/3 proteins interact with the subunits of non-canonical BAF, esBAF and npBAF complexes in pancreatic CSCs. Co-immunoprecipitation of SMAD2/3 from the nuclear lysate followed by western blotting of BRD9 and other subunits of the BAF complex. (**R**) TGFβ/Activin A signalling leads to the formation of protein complex between SMAD2/3 and BRD9/BAF that regulates the function of chemoresistance, G0 phase entry, migration and invasiveness of CSCs. Experiments represent three replicates. Statistical analysis was performed by 2-way ANOVA with multiple comparisons with Tukey correction and **** marks adjusted P-value <0.0001, *** is adjusted P-value <0.001, ** is adjusted P-value <0.01, * is adjusted P-value <0.05.

Due to the plasticity of the cell state, CSCs and non-stem cancer cells have been proposed to form a dynamic balance between differentiation and dedifferentiation. This process could be mediated by an epigenetic plasticity of the cells through the erosion of epigenetic barriers that would otherwise keep cells in a more constrained cell identity. Since we observed that BRD9 treatment abolished CSC self-renewal, we decided to study whether BRD9 inhibition could impact the re-emergence of CSC marker-expressing cells. To study this aspect, we sorted cells into CSC marker OCT4-GFP/CD133/SSEA4 negative population and analysed the re-emergence of these CSC triple marker-expressing cells upon temporary or continuous inhibition of BRD9 (**Figure 2E**). In the steady-state cell culture, the population contains approximately 1.5% OCT4-GFP+/CD133+/SSEA4+ CSCs. Control cells treated with DMSO showed the beginning of the re-emergence of OCT4-GFP+/CD133+/SSEA4+ CSCs after 3 days and reached the steady-state level similar to the untreated and sorted cells after 2 weeks. Pre-treatment of cells with I-BRD9 for 5 days significantly slowed down the re-emergence of OCT4-GFP+/CD133+/SSEA4+ CSCs, while continuous I-BRD9 treatment of cells had an even more striking effect similar to continuous treatment with TGFβ/Activin pathway inhibitor SB431542. These data suggest that BRD9 inhibition could promote CSC differentiation or decrease cellular plasticity by increasing the epigenetic barriers that are necessary for the cells to dedifferentiate from non-stem cancer cells to CSCs.

The fluorescent ubiquitination-based cell cycle indicator (FUCCI) system is a powerful tool to assess cell cycle-dependent responsiveness to drugs and the effect of drugs, gene silencing or activation on the cell cycle without the need for synchronisation^39^. We have previously used the dual-colour FUCCI system in hESCs^40^. The basic dual-colour FUCCI system uses truncated hCdt (DNA replication licensing factor) and Geminin (inhibitor of hCdt) to detect cells in G1, S and M-phases^40 41^. More recently, the presence of the cell cycle inhibitor p27 with distinct site-specific mutations has been shown to characterise G0 cells^42^. To study the effects of BRD9 epigenetic machinery inhibitors on CSC proliferation by small molecules we developed an advanced three-colour FUCCI system (RFP-hCdt1(30/120)_mAG-hGEM(1/110)_mKate2-p27(mut)) by combining the truncated hCdt1, truncated hGeminin and p27K(-). We established pancreatic ductal adenocarcinoma cell lines by TALEN-mediated targeting of the three-colour FUCCI construct into the AAV1 locus for stable genomic integration. This FUCCI system enables distinguishing between the cells in early G1, late G1, early S, S/G2/M and G0 phases (**Figure 2F-H**). We first aimed to determine the cell cycle kinetics in FUCCI cells after BRD9 inhibition. We treated the CSC-FUCCI spheres from two different PDAC cell lines for 5 days with I-BRD9, Gemcitabine or a combination of I-BRD9 and Gemcitabine, and performed flow cytometry analysis in two PDAC cell lines (**Supplemental Figure 2A-D**). In both cell lines, Gemcitabine treatment led to an increased fraction of cells positive for Cdt1-mRFP and p27k(-)-mKATE2 signals which marks G0-phase cells. Of note, the accumulated expression of p27k(-)-mKATE2 is not high in G0, which could indicate a shallow and transient quiescence with cell entry into the G0 condition while maintaining its readiness to re-enter the cell cycle phase due to KRAS activation^43^. In contrast, I-BRD9 treatment led to a reduction in G0-phase cells and also blocked cells from accumulating in the G0-phase upon Gemcitabine treatment (**Figure 2I**). This suggested that BRD9 inhibition reduces the capacity of cells to enter the G0 phase and this could be particularly useful in eliminating CSCs upon combination treatments with DNA damaging reagents such as Gemcitabine that prevent the cells from escaping genotoxic insults through temporarily dormant or quiescent states by entering the G0 phase.

PDACs are highly aggressive cancers in part due to the ability of cancer cells to migrate and initiate metastatic processes through their invasive capacities. Since BRD9 inhibition led to a reduction in CSC self-renewal and chemoresistance, we decided to investigate its impact on other key characteristics of PDAC cells such as migration capacity and invasiveness. Wound-healing assays with BRD9 genetic knockdown, chemical inhibition and protein degradation indicated reduced migration of PDAC cells compared to control cells (**Figure 2J-K**). Importantly, BRD9 knockdown also reduced PDAC cell invasiveness by trans-well assays in two separate PDAC lines (**Figure 2L-M**), indicating that BRD9 inhibition could be useful for reducing the metastatic capacity of PDACs since BRD9 inhibition decreases PDAC cell motility and invasiveness.

Collectively, chemical inhibition and knockdown of BDR9 blocked the self-renewal of CSCs, reduced CSC invasiveness and re-sensitized PDAC CSCs to conventional chemotherapeutic reagents suggesting that BRD9 could be an attractive therapeutic target in PDAC.

### BRD9/BAF complex cooperates with SMAD2/3 in regulating gene expression in pancreatic CSCs

Our previous experiments indicated that BRD9 as well as the TGF β/Activin-SMAD2/3 signalling pathway both regulate the characteristics of CSCs and PDAC cell invasiveness. This led us to hypothesize that BRD9 and TGF β/Activin-SMAD2/3 signalling pathway could cooperate in CSCs at the molecular level by protein-protein interactions. Therefore, we proceeded with identifying the binding partners of SMAD2/3 proteins in pancreatic CSCs by performing SMAD2/3 co-immunoprecipitation followed by mass-spectrometry (**Figure 2N**). This unbiased proteomic approach identified SS18/SSXT, BAF180/Polybromo-1 and ARID1A peptides indicating that these proteins could be co-factor candidates of SMAD2/3 (**Figure 2O**). STRING analysis of protein interactions confirmed that these three proteins are all part of the ATP-dependent chromatin remodelling complex BAF (BRG1/BRM-associated factor) that corresponds to the mammalian SWI/SNF complex (**Supplemental Figure 2E**). The BAF complex has tissue-specific functions that arise from the combinatorial assembly of distinct subunits such as an ES cell-specific BAF (esBAF), the neural progenitor BAF (npBAF) and neuronal BAF (nBAF), as well as cardiac progenitor-enriched (cBAF) and hematopoietic stem cell BAF^44^. More recently, a non-canonical (ncBAF, also called GBAF) complex has been described^45 46^. However, BRD9 protein is a common subunit of esBAF, npBAF, nBAF, GBAF, cBAF and the BAF complex observed in hematopoietic stem cells. Therefore, we investigated further the distinct complex composition of BAF complexes that interact with SMAD2/3 (**Supplemental Figure 2F**). SMAD2/3 co-immunoprecipitation experiments indicated an interaction of SMAD2/3 with general subunits SS18 and BRD9 (**Figure 2P**), and subunits that are specific for esBAF (BCL11a), npBAF (BAF180) and ncBAF (GLTSCR1).

These results suggest that SMAD2/3 forms a complex with several of the distinct BAF complexes (**Figure 2R**), which have all BRD9 enzyme as a subunit, and suggest that SMAD2/3 could be involved in targeting BRD9 as a subunit of the BAF complexes to distinct chromatin regions to regulate the chromatin landscape and gene expression in pancreatic CSCs.

### Inhibition of BRD9 suppresses in vivo PDAC tumour formation

To study the effect of BRD9 on PDAC cell growth *in vivo*, xenograft PDAC mouse models were developed by subcutaneous injection of CSCs from the PDAC line A13A with different transduction/treatment regimes (**Supplemental Figure 3A**). At 2 months post inoculation, only mice implanted with control adenovirus transduced cells (containing scrambled control shRNA, A13A^Null^) developed tumours. At this point, no tumours were observed in other mice implanted with Gemcitabine-treated cells (A13A^Gemcitabine^), shRNA-BRD9adenovirus-transduced cells (A13A^BRD9-KD^), or A13A^BRD9-KD^ plus with Gemcitabine treatment (A13A^BRD9-KD + Gemcitabine^), indicating the slower growth rate of these cells in contrast to A13A^Null^. After 3 months of cell inoculation, although no tumours were observed in mice with A13A^BRD9-KD + Gemcitabine^, the other mice implanted with A13A^Null^, or A13A^Gemcitabine^, or A13A^BRD9-KD^ developed tumours respectively. To further characterize these tumours, calliper measurement and PET-^18^F-FDG were performed to analyze the size and metabolic activity of these tumours. As expected, the A13A^Null^ tumours grew much faster than other tumours from month 3 to month 4, as evidenced by the significantly increased tumour volume (**Supplemental Figure 3B**). Although tumours formed by A13A^Gemcitabine^ or A13A^BRD9-KD^ were detected at month 3, they were moderate and grew slowly from month 3 to month 4. Importantly, BRD9-KD further delayed tumour growth compared with A13Al^Null^ and A13A^Gemcitabine^ groups. These results were consistent with the observation obtained from PET-^18^F-FDG. At month 4 post inoculation, the^18^F-FDG uptake in tumours formed by A13A^Null^ was much higher than in other tumours (**Supplemental Figure 3C**). Gemcitabine significantly decreased the^18^F-FDG uptake in A13A^Gemcitabine^ tumours compared with A13A^Null^ tumours, and BRD9-KD further reduced the^18^F-FDG uptake of A13A^BRD9-KD^ tumours compared with Gemcitabine treatment. Consistently, the tumours harvested from A13A^BRD9-KD^-implanted mice at month 4 were markedly smaller compared with tumours from A13A^Null^ or A13A^Gemcitabine^ implanted mice (**Supplemental Figure 3D**). The lower tumour-weight to body-weight ratios of A13A^BRD9-KD^-transplanted mice further demonstrated a progressive decrease in tumour mass (**Supplemental Figure 3E**). In contrast, no tumours were detected in any mice with A13A^BRD9-KD + Gemcitabine^ and these mice were alive without any signs of sickness at month 4. When we further extended the time window to 6 months, only one of 6 mice in the A13A^BRD9-KD + Gemcitabine^ groups developed a small tumour (data not shown). At necropsy, no metastases were observed among these tumour-bearing mice and non-tumour bearing mice. Histological examination of these tumour sections by H&E staining found prominent heterogeneity and extensive necrosis in A13A^Null^ tumours than A13A^Gemcitabine^ tumours, whereas A13A^BRD9-KD^ tumours only showed a small area of necrosis (**Supplemental Figure 3F**). These data suggested that BRD9 activity drives the tumour progression through regulating PDAC cell growth.

Although BRD9 inhibition suggested tumor progression through regulating PDAC cell growth in our cell-line xenografts, the cell-line xenografts do not accurately recapitulate the histopathological and molecular characteristics of the human parental tumor. Therefore, human xenograft PDAC models (PDX models) were employed to further evaluate the therapeutic potential of BRD9 inhibitor in vivo. Human tumors from a PDAC patient were implanted into NOD.Cg-*Prkdc*^scid^ *Il2rgtm1Wjl/*SzJ (NOD Scid gamma) host strain. Following transplantation, various treatment schedules were initiated when the tumors reached an average size of 150 mm^3^ with a repeated injection schedule (once every 3 days for a total of six treatments within 15 days) (**Figure 3A**). As expected, Gemcitabine (Gem) or combination with BRD9 inhibitor (IBRD9 + Gem) significantly delayed the tumor growth as evidenced by the tumor growth curves and tumor volume (**Figure 3B**) when compared with DMSO treated control group (Ctrl). In contrast to the Gem group the most striking tumor regressions were observed in the IBRD9 + Gem group. Notably, from day 27 to day 42 after completing treatments, the tumors in the Gem group recurred with rapid growth, whereas most of the xenograft tumors in IBRD9 + Gem group grew slowly over a prolonged period without significant changes in tumor volume (**Figure 3B**). To monitor the treatment response and tumor progression in living mice, PET-^18^F-FDG was performed (**Figure 3C**). In the contol groups, higher metabolic activity associated with higher^18^F-FDG uptake was found in these highly proliferative tumors. In contrast, treatment with either Gem or IBRD9 + Gem decreased levels of tumor^18^F-FDG uptake. In particular,^18^F-FDG uptake in the IBRD9 + Gem group was significantly lower than in any other group (**Figure 3C**) indicating that the combination of IBRD9 + Gem significantly delayed the tumor growth. Consistent with reduced tumor growth, the tumor size and tumor-weight to body-weight ratios in IBRD9 + Gem group were significantly lower than those originating from Ctrl and Gem treatment groups (**Figure 3D-E**). Importantly, the combination therapy with IBRD9 did not show any toxic effects as no significant alteration of body weight was observed in IBRD9 + Gem treated mice (data not shown). Histology analysis of these tumor sections revealed a larger necrosis area in Ctrl and Gem groups as compared with IBRD9 + Gem groups (**Figure 3F**), indicating rapid growth of these tumors in Ctrl and Gem groups. In line with these findings, Ki67^+^ proliferating tumor cells were substantially increased in Ctrl and Gem groups, whereas they were reduced in IBRD9 + Gem treated tumors (**Figure 3G-H**), suggesting a potential additive effect of BRD9 in controlling primary tumor growth.

**Figure 3.**
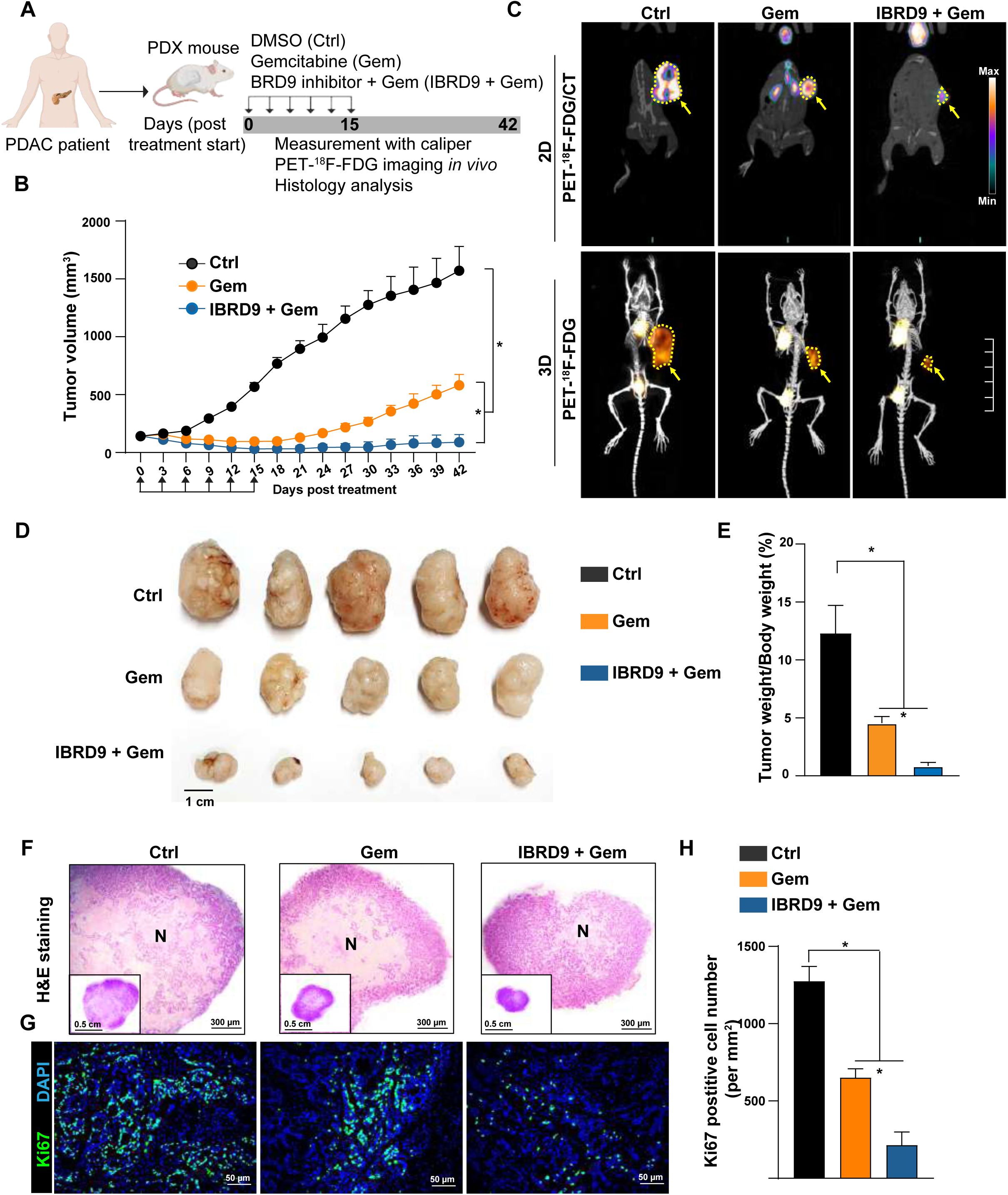
Inhibition of BRD9 enhances antitumor activity in patient-derived xenograft (PDX) models of PDAC. (**A**) Experiment scheme for assessing BRD9 inhibitor in PDX models. (**B**) Tumor growth in PDX models with various treatment (n=5). Statistical analysis was performed by two-way ANOVA with multiple comparisons with Tukey post-test. *P<0.05. (**C**) Representative images of PET-^18^F-FDG /computed tomography scanning. (**D**) Gross morphology of tumors harvested from PDX mouse models. (**E**) Measurement of tumor-weight to body-weight ratio (n=5). Statistical analysis was performed by one-way ANOVA with Dunnett’s post-test. *P<0.05. (**F**) Representative images of H&E stained tumor sections obtained from PDX mouse models. N denotes tumor necrosis. (**G**) Representative images of immunofluorescence staining with Ki67 antibody in tumor sections obtained from PDX mouse models. (**H**) Quantification data of Ki67 immunofluorescence staining (n=5). Statistical analysis was performed by one-way ANOVA with Dunnett’s post-test. *P<0.05.

### BRD9 inhibition eliminates CSCs from patient tumour

CSCs are particularly challenging to eliminate due to their chemoresistant phenotype. Our experiments so far showed that treatment of PDAC cells with BRD9 inhibitors reduced the presence of CSCs while this effect was further enhanced by the combined treatment of BRD9 inhibitors and Gemcitabine. Although encouraging, these results were observed in PDAC cell lines while cells from primary pancreatic tumours could respond differently. To further examine the translational relevance of BRD9 as a candidate therapeutic target, we utilized freshly isolated primary cancer patient samples and treated them with I-BRD9 or in combination with Gemcitabine for 72h followed by single-cell RNA-sequencing (**Figure 4A**) for ∼12,000 cells. The analysis of this scRNA-sequencing data indicated different populations of cells based on their expression of genes characteristic for ductal cells (*KRT19*), fibroblasts (*LUM*) and stellate cells (*THY1*), and the further separation of pancreatic ductal cells to different sub-population of cancer cells (**Figure 4B**, **D**). One of these ductal cell subpopulations, designated as Ductal cells 3, expressed a range of cancer stem cell markers. This cancer cell subpopulation was reduced by I-BRD9 treatment (from 19.4% to 6.8%) and showed a ten-fold reduction in numbers (from 19.4% to 1.9%) upon co-treatment with I-BRD9 and Gemcitabine (**Figure 4C**). Enriched Gene Ontology terms for downregulated genes in conditions I-BRD9/Gemcitabine and I-BRD9 versus DMSO and Gemcitabine indicated central mechanisms that are relevant for PDAC development and CSCs. These terms included Extracellular matrix organization, cell proliferation, cell migration, cell adhesion, regulation of apoptotic signalling pathways, and regulation of angiogenesis (**Supplemental Figure 4A**), all of particularly relevance for tumour aggressiveness and metastatic processes. The enriched Pathways for downregulated genes in these same conditions uncovered TGF β/Activin A signalling pathway, fibrosis, VEGFA signalling, wound healing, autophagy pathways in cancer, proinflammatory/profibrotic mediators, NFκB and YAP1/ECM axis (**Supplemental Figure 4B**). Among the genes that are specifically repressed by I-BRD9 treatment in the CSC population of cells were *SOX4* and *TWIST1* (**Figure 4E**), and *SNAIL2*, *JUN*, *IL6* and *IGF1* (**Supplemental Figure 4C**). The combined treatment of I-BRD9 and Gemcitabine resulted in a synergistic reduction of these CSC factors (**Figure 4E** and **Supplemental Figure 4C**). Altogether, the results using resected patient tumour samples indicated that BRD9 inhibition by a small molecule compound can efficiently target and eliminate the CSC subpopulation of pancreatic cancer cells, thus confirming our prior discoveries on PDAC cell lines.

**Figure 4.**
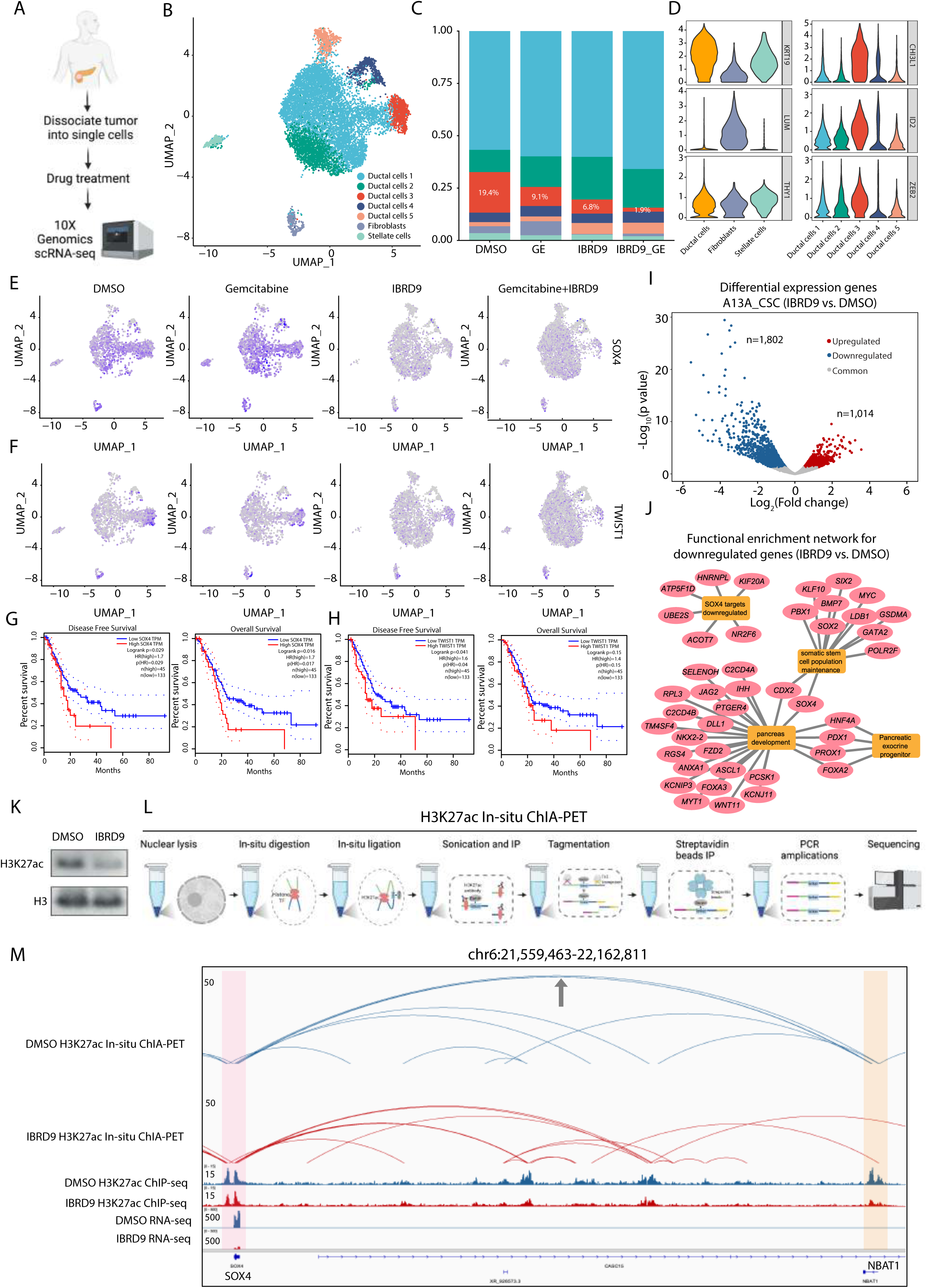
Inhibition of BRD9 reduces CSCs in patient tumors and the enhancer-promoter connectome in CSCs. (**A**) Schematic depiction of single-cell RNA-sequencing strategy. The surgically removed PDAC tumour was dissociated to single cells and divided equally into four treatment conditions, control, Gemcitabine, I-BRD9 and Gemcitabine + I-BRD9. After 72 hours the samples were processed for single-cell RNA-sequencing. (**B**) Single-cell RNA-sequencing analyses of patient derived primary PDAC cells with clustering of cells. (**C**) BRD9 inhibition reduces the relative abundance of stem cell-like cells and enhances the effects of Gemcitabine. (**D**) Violin plots corresponding to marker gene expression in the different cell types in clusters. (**E-F**) BRD9 inhibition leads to the elimination of cells that express CSC genes (**E**) *SOX4* and (**F**) *TWIST1*. Expression levels of *SOX4* and *TWIST1* in the CSC population is shown as dot plot graphs where blue colour indicates higher expression level compared to gray colour. (**G-H**) Higher expression of *SOX4* and *TWIST1* correlate with lower survival of pancreatic cancer patients, and lower disease free survival based on TCGA data. (**I**) Differential gene expression analysis upon BRD9 inhibition in CSCs. RNA-sequencing identified 1,802 downregulated genes and 1,014 upregulated genes in I-BRD9 treated samples compared to control cells. (**J**) Functional enrichment network of downregulated genes upon BRD9 inhibition uncover factors regulating stem cell maintenance, pancreatic development and pancreatic exocrine progenitors, as well as SOX4 target genes. (**K**) BRD9 inhibition reduces the abundance of H3K27ac in CSCs. Western blot analysis of the H3K27ac in control and I-BRD9 treated samples while total Histone H3 acts as a loading control. (**L**) Schematic depiction of analysing enhancer-promoter connectivity by H3K27ac in-situ ChIA-PET, H3K27ac ChIP-seq and RNA-seq. (**M**) Genome browser view of H3K27ac loops for *SOX4* genes. The top 2 rows represent the H3K27ac HiChIP loops, middle 2 rows represent H3K27ac ChIP-seq and the bottom 2 rows represent RNA-seq in Ctrl and IBRD9 treated A13A CSCs. *SOX4* promoter is highlighted in pink and *NBAT1* promoter (enhancer-like) is highlighted in orange.

### BRD9 inhibition disrupts enhancer-promoter connectome of stemness-related genes in pancreatic CSCs

Next, we performed bulk RNA-sequencing upon I-BRD9 inhibition in A13A CSCs which identified 1,802 downregulated genes and 1,014 upregulated genes (**Figure 4I**). The functional enrichment network analysis of the downregulated genes by BRD9 inhibition revealed a number of central transcriptional regulators of somatic stem cell maintenance (e.g. *SOX4*, *SOX2*, *MYC*, *BMP7*), pancreatic exocrine progenitors/pancreatic development (e.g. *DLL1*, *JAG2*, *WNT11*, *NKX2-2*) and SOX4 target genes (**Figure 4J**). GSEA pathway analysis of I-BRD9 downregulated genes indicated SMAD3 mediated transcription and Epithelial-to-Mesenchymal Transition (**Supplemental Figure 4A**) as enriched pathways. Altogether these results emphasize the importance of key processes involved in regulating stem cell maintenance and metastasis that are downregulated upon BRD9 inhibition.

In a cellular context, transcriptional output is largely orchestrated by cis-regulatory elements (CREs), in particular enhancers and promoters. By performing ATAC-seq assay, we identified that I-BRD9 treatment in A13A CSCs results in extensive loss of CREs (n=1,609) compared to control samples (**Supplemental Figure 4D**). In line with this, Western blots demonstrated that acetylated lysine27 at histone 3 (H3K27ac)^47^, an important active enhancer and promoter histone mark, was significantly decreased in response to BRD9 inhibition (**Figure 4K**). In most cases, enhancers control gene expression through long-range interactions with promoters^48 49^, but very little is known about the enhancer/promoter connectome in pancreatic CSCs. In order to study the effects of BRD9 inhibition on the enhancer/promoter connectome of CSCs, we performed H3K27ac In-situ ChIA-PET^50-52^ experiments, which capture H3K27ac-centric chromatin interactions (i.e. enhancer/promoter connectome) (**Figure 4L**). These data suggest that the transcriptional downregulation of stemness-related genes is possibly due to the loss of long-range enhancer-promoter connectome. For example, we found that in Control CSCs, the *NBAT1* promoter is characterized by H3K27ac enrichment on the promoter but is transcriptionally silenced. In addition, the *NBAT1* promoter has strong chromatin interaction with the *SOX4* promoter, suggesting that this non-transcribing promoter may function as an enhancer^53^, to regulate the expression of SOX4. However, I-BRD9 treatment diminished the H3K27ac enrichment on the *NBAT1* promoter and its physical contact with the *SOX4* promoter, which possibly contributes to the transcriptional downregulation of *SOX4* (**Figure 4M**). Similarly, the transcriptional downregulation of other stemness-related genes (e.g. *CD133/PROM1*, *SMAD1* and *SNAI2*) are linked to the loss of enhancer-promoter connectome (**Supplemental Figure 5B**). ATAC-seq data showed SMAD2/3 footprints at the anchor regions (**Supplemental Figure 5C-D**). SMAD3 has several TF motifs in the JASPAR motif database. We identified MA0513 in Ctrl-specific ATAC peaks (254 hits) and 211 hits in IBRD9-specific ATAC peaks, while MA1622 has 1370 hits in total on anchor regions with approximately similar number of hits in both conditions. Hence, SMAD2/3 footprints were not significantly lost upon BRD9 inhibition. To further study the binding dependency of BRD9 and SMAD2/3 on TGF β/Activin pathway we performed BRD9 and SMAD2/3 ChIP-qPCR on representative SMAD2/3 binding regions near SOX4, CD133/PROM1, SNAI2 and SMAD1 loci upon TGF β/Activin pathway inhibitor SB431542 treatment of pancreatic CSCs (**Supplemental Figure 5E**). BRD9 and SMAD2/3 bound to the anchor regions, while the binding of BRD9 on the anchor regions Β/Activin pathway inhibition. These results indicate that BRD9 and SMAD2/3 are co-binding to these regions and SMAD2/3 transcription factors recruit BRD9 to the gene promoter and enhancer regions (**Supplemental Figure 5F**). Collectively, our results revealed the cooperation of BRD9 and SMAD2/3 in regulating the enhancer-promoter connectome and gene expression of stemness-related genes in pancreatic CSCs.

## Discussion

Epigenetic regulation of gene expression is essential for guiding developmental processes throughout embryogenesis and tissue homeostasis in an adult organism. The cellular identity of differentiated somatic cells can also be epigenetically reprogrammed through erasure and rewriting of chromatin marks resulting in a pluripotent stem cell state. Similar differentiation and dedifferentiation processes mediated by transcription factors and epigenetic regulatory proteins can contribute to tumour formation, invasiveness, metastatic processes, dormancy and reactivation of cancer cells, clonal evolution of tumour cells and the development of therapeutic resistance in cancers^4^. Of particular importance to all these processes, epigenetic mechanisms regulate phenotypic plasticity and the self-renewal capacity of cancer stem cells. Accordingly, targeting epigenetic mechanisms offers an attractive strategy to eliminate cancer stem cells through reducing their self-renewal characteristics or re-sensitizing them for chemotherapeutic drugs.

We used a compound screening approach to identify candidate targets that regulate stem cell-like characteristics of pancreatic cancer stem cells. The compound library used consisted of extensively ex vivo validated small molecule compounds covering major epigenetic target classes. For several of the targets, including BRD9, the inclusion of multiple chemical scaffolds for a given target and availability of negative controls (similar chemical scaffolds with significantly reduced activity) provided a robust approach for target identification. Using this strategy we identified BRD9 as a novel epigenetic regulator of pancreatic CSC and the corresponding small molecule inhibitors for this enzyme. Our experiments uncovered that BRD9 as a subunit of ncBAF, esBAF and npBAF cooperates with TGF β/Activin-SMAD2/3 in regulating pancreatic CSC self-renewal, chemoresistance and invasiveness. SMAD2/3 recruit the BAF complexes to the expression of stem cell loci by regulating the 3D enhancer and promoter interactions in CSCs. This provides important mechanistic insight to the transcriptional complexes that target epigenetic regulators and identifies the epigenetic regulatory complexes that are linked to TGF β/Activin-SMAD2/3 signalling pathway in PDACs. TGF β/Activin-SMAD2/3 signalling pathway has been known to play a dual function in tumorigenesis by exhibiting anti-tumorigenic function in early stages of tumorigenesis and pro-tumorigenic function in later stages of tumorigenesis. Our results provide particular insight to the role of TGF β/Activin-SMAD2/3 in promoting CSC characteristics by directly cooperating with the BAF complexes in regulating stemness circuitries. The fact that multiple BAF complexes are present in pancreatic CSCs is interesting given that each BAF complex is generally associated with distinct stem cell types. esBAF functions in regulating the self-renewal and pluripotency of embryonic stem cells, whereas npBAF is important for neural progenitor/neural stem cells^60^, and non-canonical BAF is linked to naïve pluripotent stem cells^46^. This could reflect the developmentally plastic state of pancreatic CSCs and their capacity to reversibly differentiate and de-differentiate by using BAF complexes normally regulating the naïve ESC, primed ESC and neural stem cell/neural progenitor cellular identities. The similarity of some pathways in pancreatic cancer with neural development during embryogenesis has been also indicated by the presence of mutations in axon guidance pathways in PDACs. SMAD2/3 binds several of the BAF complexes in CSCs: ncBAF, esBAF and npBAF. Hence, the dynamics of different SMAD2/3-BAF complexes could provide the necessary stem cell-like developmentally plastic or “metastable” capacity of CSCs. We have previously shown that TGF β/Activin-SMAD2/3 regulates pluripotency in human embryonic stem cells through MLL/COMPASS mediated histone modifications^24^. Based on our current data it seems that TGF β/Activin-SMAD2/3 also promotes the stem cell-like characteristics of pancreatic CSC. In the latter case, SMAD2/3 seem to direct the BAF complex to stem cell loci to induce their expression.

Many cancers have been found to be almost separate diseases due to different mutations and epigenetic effects that impact divergent mechanisms during tumorigenesis. This heterogeneity in the disease has been described most comprehensively in breast cancer while non-genomic mechanisms and genomic mutations also cause significant heterogeneity in PDACs. Since BAF subunits ARID1A and BAF180/Polybromo are mutated in a subset of PDACs, it suggests possibly divergent mechanisms how the BAF complexes could promote CSCs characteristics in PDACs. Furthermore, SMAD4 as an important cofactor of SMAD2/3 and component of TGF β/Activin signalling pathway is mutated in approximately half of PDACs^22^. Therefore, it will be important to determine if the absence of SMAD4 can impact the function of BAF complexes in pancreatic CSCs and whether BRD9 inhibition is also an attractive therapeutic target in SMAD4 mutated PDACs or only in SMAD4 wild-type PDACs. This could indicate differences in the molecular mechanisms of how CSCs are epigenetically regulated upon SMAD4 mutation. How it impacts disease progression and patient survival needs further research.

An interesting characteristic of tissue-specific stem cells is their capacity to enter and exit a quiescent stage. A similar process is believed to occur for CSCs during which the cancer cell is presumed to enter a dormant stage in the G0 phase. This is intriguing, considering that a large majority of PDACs have early activating mutations in KRAS turning it into a constitutively activated oncogene that promotes cell proliferation. By developing a novel tool (3FUCCI) for cell cycle analyses in pancreatic CSCs we were able to investigate the cell cycle progression of CSCs and the effects of BRD9 on cell cycle. Our experiments revealed that BRD9 inhibition prevents CSCs from entering the G0 phase. This inability to slow down or exit the proliferative cycle in G1 and enter the quiescent state G0, makes the cells susceptible for Gemcitabine-mediated killing. Interestingly, the G0 phase in the PDAC CSCs has only a low p27(K-) induction. This could reflect that cells are temporarily in a ‘‘shallow quiescent’’ state, which has been termed ‘‘GAlert’’ or ‘‘primed’’ cells. Such cells can re-enter G1 and the cell cycle faster than more ‘‘deeply quiescent’’ cells. Signalling through mTORC1 has been shown to control the transition from G0 to GAlert in Muscle Stem Cells (MuSCs), whereas the primed state in Hematopoietic Stem Cells (HSCs) has been shown to be regulated by the level of CDK6^43^. Future experiments should further investigate the mechanisms how BRD9 can regulate the entry of PDAC cells into the G0 phase and whether this is coordinated with the mTORC1 pathway and CDK6.

Our research validated the translational relevancy of our preclinical models by using surgically removed tumor samples from PDAC patients. The BAF complex with the BRD9 epigenetic modulator serves as novel, first-in-class anti-cancer compound that is expected to enter clinical testing in the future. This novel approach is particularly attractive in pancreatic cancer which is in urgent need for better selection of targets, in particular targeting the heterogenous cancer cell populations including chemoresistant CSCs. BRD9 inhibition could be used in combination with currently available treatments such as Gemcitabine or FOLFIRINOX with the objective of re-sensitizing pancreatic CSCs that are particularly resistant to currently used chemotherapies. Hence, the improved molecular target identification serves a dual purpose; in addition to improving drug selectivity, our discoveries will guide effective therapy combination with either standard of care or other investigational agents. The determination of the CSC 3D chromatin architecture provides insights into higher levels of epigenetic regulation of gene expression. Our results indicated the SMAD2/3-BAF complex regulates promoter-enhancer connections in pancreatic CSCs and upon BRD9 inhibition, enhancer-promoter looping at various cancer stemness genes such as SOX4 are lost, thus leading to stem cell gene repression and loss of CSC self-renewal. Targeting the regulators of enhancer-promoter connectomes is thus emerging as an attractive therapeutic strategy for eliminating the stem cell-like populations in PDACs. Our findings can potentially translate into clinical benefit for PDAC patients that have a surgically unresectable cancer and even undergoing relapse.

## Acknowledgements

This work was supported by a Cancer Research UK Career Development Fellowship (Grant ID C59392/A25064 to Dr Pauklin) and Pancreatic Cancer UK (Grant/Award Number: 2018RIF_03 to Dr Pauklin); the National Natural Science Foundation of China (grants 81800262 to Dr Jiang); Natural Science Foundation of Guangdong Province (grant 2018A030313029 to Dr Jiang); Science and Technology Planning Project of Guangzhou (grant 201903010005 to Dr Jiang); National Institutes of Health (R01 HL157456 and R01 HL143490 to Dr Y. Wang); Clarendon Fund and St Edmund Hall Scholarship (grant SFF1920_CB_MSD_759707 to Dr Feng); The Kennedy Trust for Rheumatology Research (Daphne Jackson Fellowship to Dr Chang); Kristjan Jaak Scholarship Programme (Ms Ervin); Cancer Research UK DPhil in Cancer Science grant (Ms Wu); China Scholarship Council – University of Oxford Scholarship, (SFF2122_CSCUO_1284663 to Mr Deng (No. 202108330024)); National Institutes of Health (grants R01 AR078001, R01 HL130356, R01 HL105826, R38 HL155775, and R01 HL143490 to Dr Sadayappan).

## Materials and Methods

For full materials and methods please see Supplemental Information.

### The small molecule screening library

The screening library contained concentrated small molecule compounds with verified biochemical activity against their targets. Most of the compounds target epigenetic regulators with high specificity (**Supplemental Table 1**).

### Cell lines and cell culture

The A13A, A13D, A13B cells were provided by Christine Iacobuzio-Donahue at The Memorial Sloan Kettering Cancer Center. FG and L3.6 were provided by Isaiah J. Fidler at The University of Texas MD Anderson Cancer Center. For standard cell cultures, cells were grown at 37°C humidified incubator containing 5% CO2 in Dulbeccos modified Eagles medium (DMEM), high glucose, GlutaMAX™ Supplement, pyruvate (Thermo Fisher Scientific, Inc.), 100 U/ml penicillin, 100 mg/ml streptomycin (Thermo Fisher Scientific, Inc.), MEM Non-Essential Amino Acids Solution 1X (Life Technologies; Thermo Fisher Scientific), MEM Vitamin Solution 1X (Life Technologies; Thermo Fisher Scientific) and 10% inactivated fetal calf serum (FCS; Life Technologies). For 3D cultures, cells we grown in ultra-low attachment plates (Thermo Fisher Scientific, Inc.) in Dulbecco’s Modified Eagle Medium/F12 (Sigma-Aldrich) supplemented with 2.5mM L-Glutamine (Thermo Fisher Scientific, Inc.), 1 % B27 supplement (Life Technologies), 100 U/ml penicillin, 100 mg/ml streptomycin (Thermo Fisher Scientific, Inc.), 20ng/ml basic thermostable fibroblast growth factor (FGF-Basic TS, Proteintech) (5,000 cells/ml).

### Tumour sphere assays

Cells are seeded into ultra-low attachment plates in Dulbecco’s Modified Eagle Medium/F12 (Sigma-Aldrich) supplemented with 2.5mM L-Glutamine (Thermo Fisher Scientific, Inc.), 1 % B27 supplement (Life Technologies), 100 U/ml penicillin, 100 mg/ml streptomycin (Thermo Fisher Scientific, Inc.), 20ng/ml basic thermostable fibroblast growth factor (FGF-Basic TS, Proteintech) (5,000 cells/ml). For tumour sphere formation assay, cells were passaged after one-week incubation, and grown for another week after which the tumour sphere numbers were counted under a phase-contrast microscope using the 40x magnification lens or by Celigo Image Cytometer (Nexcelom). For RNA-seq and ATAC-seq experiments, serial passaging of the first-generation tumour spheres is required. After 7 days of incubation the tumour spheres were harvested by using a 40 µm cell strainer and centrifuged for 5 min at 200 x g at RT. Dissociate the pellet of tumour spheres to single cells using trypsin, and then expanded for another 4 days before performing the treatments of samples.

## Materials and methods

### Knockin cell lines for Oct4

pCCC construct and OCT4 TALEN constructs (pTALEN_V2-OCT4F, pTALEN_V2-OCT4R) constructs were a gift from Francis C. Lynn and have been published^1^. OCT4-eGFP-PGK-Puro was a gift from Rudolf Jaenisch (Addgene plasmid #31937) and have been published^2^. Cells were transfected with Lipofectamine 3000 (ThermoFischer Scientific) and cultured for 4 days after transfection before selecting with 0.25 µg/mL puromycin (Sigma). Colonies were individually picked, trypsinized and placed into 24-well plates with 500 µl of media. Once clones were close to confluent, cells were replica plated for genotyping, freezing and for expanding the correctly targeted clones. Genomic DNA was extracted using Promega Wizard SV Genomic DNA Purification System (Promega) and genotyping was performed as described^1^. Positive clones were analysed by flow cytometry to estimate the frequency of eGFP positive cells in the cancer cell population. We additionally used primers GGTGCTCAGGTAGTGGTTGTCG and CTCTAATGTCCTCCTCTAACTGCTCTAGG for Oct4-GFP region verification, as well as CACAACCTCCCCTTCTACGAGC and GCATCATTGAACTTCACCTTCCCTC for Oct4-Puromycin region verification.

### The small molecule screening library

The screening library contained concentrated small molecule compounds with verified biochemical activity against their targets. Most of the compounds target epigenetic regulators with high specificity (Supplementary Table 1).

**Supplementary Table 1.** Small molecule compounds used in the screening experiment. List with compound names, working concentrations and target molecules.

| Vial | Source Name | [Working]<br>uM | Class/Target |
| --- | --- | --- | --- |
| 1 | (+)-JQ1 | 1 | Bromodomains - BRD2, BRD3, BRD4, BRDT (BET) |
| 2 | (-)-JQ1 (inactive) | 1 | Bromodomains - Negative control |
| 3 | PFI-1 | 5 | Bromodomains - BRD2, BRD3, BRD4, BRDT (BET) |
| 4 | I-BET | 1 | Bromodomains - BRD2/3/4 |
| 5 | Bromosporine | 1 | Bromodomains - pan-Bromodomain |
| 6 | CBP/BRD4 (0383) | 5 | Bromodomains - CBP, BRD4(1) |
| 9 | SGC-CBP30 | 1 | Bromodomains - CREBBP, EP300 |
| 10 | I-CBP112 | 1 | Bromodomains - CREBBP, EP300 |
| 15 | RVX-208 | 5 | Bromodomains - BRD2, BRD3, BRD4, BRDT (BET, BD2) |
| 16 | SMARCA | 2.5 | Bromodomains - SMARCA, PB1 |
| 17 | PB1/SMARCA | 1 | Bromodomains - SMARCA, PB1 |
| 18 | PFI-3 | 1 | Bromodomains - SMARCA2/4, PB1(5) |
| 19 | GSK2801 | 1 | Bromodomains - BAZ2A, BAZ2B |
| 20 | PFI-4 | 1 | Bromodomains - BRPF1B |
| 21 | TRIM24/BRPF | 10 | Bromodomains - TRIM24/BRPF |
| 22 | OF-1 | 5 | Bromodomains - pan-BRPF |
| 23 | Belinostat | 5 | HDAC - hydroxamic acids |
| 24 | CXD101 | 1 | HDAC - |
| 25 | Valproic acid | 1000 | HDAC - aliphatic acid compounds |
| 26 | Entinostat | 0.5 | HDAC - ortho-amino anilides |
| 27 | SAHA | 2.5 | HDAC - hydroxamic acids |
| 28 | Trichostatin A | 0.5 | HDAC - hydroxamic acids - Class I & II |
| 29 | SRT1720 | 1 | HDAC - SIRT1 (indirect?) activator |
| 30 | EX 527 | 1 | HDAC - SIRT1 |
| 32 | CI-994 | 1 | HDAC - 1,2,3,(8) |
| 33 | CPI-360 | 10 | Histone methyltransferase - EZH2 and EZH1 |
| 34 | CPI-413 | 10 | Histone methyltransferase - EZH2 and EZH1 |
| 35 | UNC0638 | 1 | Histone methyltransferase - G9a, GLP |
| 37 | UNC0642 | 1 | Histone methyltransferase - G9a, GLP |
| 38 | A-366 | 2 | Histone methyltransferase - G9a, GLP |
| 39 | Chaetocin | 0.05 | Histone methyltransferase - SUV39H1 |
| 43 | PFI-2 | 2 | Histone methyltransferase - SETD7 |
| 45 | SGC0946 | 7.5 | Histone methyltransferase - DOT1L |
| 46 | GSK343 | 3 | Histone methyltransferase - EZH2 |
| 47 | UNC1999 | 1 | Histone methyltransferase - EZH2 |
| 49 | LLY-507 | 1 | Histone methyltransferase - SMYD2 |
| 50 | Tranylcypromine | 20 | Lysine demethylases - LSD1 |
| 51 | GSK-LSD1 (irreversible) | 0.5 | Lysine demethylases - LSD1 |
| 52 | GSK690 | 5 | Lysine demethylases - LSD1 |
| 53 | GSK J4 | 10 | Lysine demethylases - JMJD3, UTX, JARID1B |
| 54 | GSK J5 (inactive) | 10 | Lysine demethylases - Negative control |
| 55 | IOX1 (5-carboxy-8HQ) | 40 | Lysine demethylases - pan-2-OG |
| 57 | Methylstat (Ester) | 2.5 | Histone demethylase |
| 58 | (E)-JIB-04 | 0.05 | Histone demethylase - Pan JmjC |
| 60 | ML324 | 5 | Histone demethylase - JMJD2E |
| 61 | IOX2 | 10 | Prolyl-Hydroxylases - PHD2 (EGLN1) |
| 62 | OICR-9429 | 1 | Methyl Lysine Binder? - WDR5 |
| 63 | UNC1215 | 5 | Methyl Lysine Binder - L3MBTL3 |
| 64 | 5-Azacitidine | 10 | DNA methyltransferase (DNMT) - |
| 65 | 5-Azadeoxycytidine | 5 | DNA methyltransferase (DNMT) - DNMT1/3 |
| 66 | Olaparib | 1 | Poly ADP ribose polymerase (PARP) |
| 67 | Rucaparib | 10 | Poly ADP ribose polymerase (PARP) |
| 68 | K00135 | 1 | Kinase inhibitor - ATP competitive - PIM |
| 69 | 5-Iodotubercidin | 1 | Kinase inhibitor - ATP mimetic - Haspin |
| 70 | C646 | 1 | Histone acetyltransferase (HAT) p300/CBP |
| 74 | DUAL1946 | 1 |  |
| 75 | GSK484 | 1 | Peptidyl arginine deiminase (PAD4) |
| 76 | KDOBA67 | 10 | Histone demethylase |
| 77 | BAZ2-ICR | 1 | Bromodomains - BAZ2A, BAZ2B |
| 78 | NI-57 | 1 | Bromodomains - pan-BRPF |
| 79 | LP99 | 1 | Bromodomains - BRD9, BRD7 |
| 80 | SGC707 | 1 | Arginine methyltransferase - PRMT3 |
| 81 | RGFP966 | 10 | HDAC - HDAC3 |
| 82 | PCI-34051 | 5 | HDAC - HDAC8 |
| 83 | Rocilinostat | 10 | HDAC - HDAC6 |
| 84 | Tubastatin A HCl | 10 | HDAC - HDAC6 |
| 85 | KDOAM-25a | 1 | Lysine demethylases - JARID |
| 86 | KDM5-C70 | 10 | Histone demethylase - JARID1 |
| 88 | MAZ1805 | 1 | t-RNA sythetase |
| 89 | MAZ1392 | 1 | t-RNA sythetase |
| 90 | BI-9564 | 1 | Bromodomains - BRD9, BRD7 |
| 91 | NVS-CECR2-1 | 1 | Bromodomains - CECR2 |
| 92 | GSK106 | 1 | Peptidyl arginine deiminase (PAD4) |
| 93 | J556-42R | 1 | Arginine methyltransferase - PRMT5 |
| 94 | J556-63R | 1 | Arginine methyltransferase - PRMT5 |
| 95 | J556-70R | 1 | Arginine methyltransferase - PRMT5 |
| 96 | A-196 | 1 | Histone methyltransferase - SUV420H1/H2 |
| 97 | BAY-598 | 1 | Histone methyltransferase - SMYD2 |
| 98 | J556-143 | 1 | Arginine methyltransferase - PRMT5 |
| 99 | MS049 | 1 | Arginine methyltransferase |
| 100 | MS023 | 1 | Arginine methyltransferase - Type I PRMTs |
| 101 | MS003 | 1 | Arginine methyltransferase - negative control |
| 104 | SGI-1776 | 10 | Kinase inhibitor - Haspin |
| 105 | CHR-6494 | 1 | Kinase inhibitor - Haspin |
| 106 | CPI-169 | 10 | Histone methyltransferase - EZH2, EZH1 |
| 107 | UNC2400 | 1 | Histone methyltransferase - EZH2 |
| 108 | GSK864 | 5 | Dehydrogenase |
| 109 | GSK8814 | 10 | Bromodomains - ATAD2 |
| 110 | GSK8815 | 10 | Bromodomains - ATAD2 |
| 111 | GSK959 | 1 | Bromodomains - BRPF1 |
| 112 | NVS-CECR2-C | 1 | Bromodomains - CECR2 |
| 113 | BAY-299 | 1 | Bromodomains - BRD1, TAF1 |
| 114 | PCI-24781 | 10 | HDAC - |
| 115 | Romidepsin | 1 | HDAC - |
| 116 | Mocetinostat | 10 | HDAC - |
| 117 | Santacruzamate | 50 | HDAC 2? |
| 118 | KDOAM32 | 1 | Lysine demethylases - JARID |
| 120 | MS409N | 1 | Arginine methyltransferase - PRMT4, PRMT6 inactive control |
| 121 | TP-064 | 1 | Arginine methyltransferase - PRMT4 |
| 122 | TP-064N | 1 | Arginine methyltransferase - PRMT4 |
| 123 | A-395 | 1 | Methyl Lysine Binder - EED |
| 124 | A-395N | 1 | Methyl Lysine Binder - EED |
| 125 | I-BRD9 | 10 | Bromodomains - BRD9 |
| 126 | TP-472 | 1 | Bromodomains - BRD9 |
| 127 | TP-472N | 1 | Bromodomains - BRD9 |
| 128 | KDOPZ-32a | 1 | Lysine demethylases - KDM5 |
| 129 | KDOOA012000 | 1 | Lysine demethylases KDM2 |
| 130 | AMI-1 | 50 | Arginine methyltransferase - PRMT |
| 131 | TMP269 | 10 | HDAC -4, 5, 7 &9 |
| 132 | AGK2 | 10 | HDAC - SIRT2 |
| 133 | GSK6853 | 1 | Bromodomains - BRPF1/2/3 |
| 134 | GSK9311 | 1 | Bromodomains - BRPF1/2/3 |
| 135 | LLY-283 | 1 | Arginine methyltransferase - PRMT5 |
| 136 | TD001851a | 1 | Methyl Lysine Binder/tudor domain -Spin1 |
| 137 | TDOSI000058a | 1 | Methyl Lysine Binder/tudor domain -Spin1 |
| 138 | TD001863a | 1 | Methyl Lysine Binder/tudor domain -Spin1 |
| 139 | TDOSI000062a | 1 | Methyl Lysine Binder/tudor domain -Spin1 |
| 140 | TD001857a | 1 | Methyl Lysine Binder/tudor domain -Spin1 |
| 141 | TD001856a | 1 | Methyl Lysine Binder/tudor domain -Spin1 |
| 142 | TD001858a | 1 | Methyl Lysine Binder/tudor domain -Spin1 |
| 143 | TMP195 | 1 | HDAC -4,5,7,9 |
| 144 | GSK2879552 | 10 | Lysine demethylases - LSD1 |
| 145 | TDO20821a | 1 | Methyl Lysine Binder/tudor domain -Spin1 |
| 146 | TDO20824a | 1 | Methyl Lysine Binder/tudor domain -Spin1 |
| 147 | TDO20823a | 1 | Methyl Lysine Binder/tudor domain -Spin1 |
| 148 | A-485 | 1 | Histone acetyltransferase (HAT) p300/CBP |
| 149 | A-486 | 1 | Histone acetyltransferase (HAT) p300/CBP |
| 150 | GSK4027 | 1 | Bromodomains - PCAF, GCN5 |
| 151 | GSK4028 | 1 | Bromodomains - PCAF, GCN5 |
| 152 | L-Moses | 1 | Bromodomains - PCAF, GCN5 |
| 153 | D-Moses | 1 | Bromodomains - PCAF, GCN5 |
| 154 | PFI-5 | 1 | Histone methyltransferase - SMYD2 |
| 155 | YX39-31b | 1 | Methyl Lysine Binder/tudor domain -Spin1 |
| 156 | TDO208229 | 1 | Methyl Lysine Binder/tudor domain -Spin1 |
| 157 | TDO01856a | 1 | Methyl Lysine Binder/tudor domain -Spin1 |
| 158 | TDO20826a | 1 | Methyl Lysine Binder/tudor domain -Spin1 |
| 159 | Bortezomib | 0.1 | Proteasome |
| 160 | Carfilzomib | 0.1 | Proteasome |
| 161 | RTS-V5 | 1 | Proteasome and HDAC |
| 162 | dBRD9 | 1 | Bromodomains - BRD9 |
| 163 | BI-7273 | 0.1 | Bromodomains - BRD9/7 (IC50 19/117nM) |
| 164 | CPI-621 | 0.1 | Lysine demethylases - KDM5 |

### Screening of the chemical compounds

The cells were grown in 96-well plates in standard growth medium with puromycin (1 µg/ml stock). Three technical replicates and three biological replicates were used for the screening. Cells were plated at a concentration of 10,000 cells in 100 µl of media per well in a 96-well plate. One day after plating the cells, the medium was changed to 90 µl standard growth medium supplemented with puromycin (0.5 µg/ml) and Activin A (10 ng/ml). On the same day, the compounds were added: first, 100x compound library dilutions were made, and 10 µL of 100x diluted chemical was added to each well to obtain 1000x final dilution of the compounds. Cells were then cultured with chemical compounds for five days with media change at day 0, day 2 and day 4 supplemented by fresh compounds. Each replicate was analyzed using Celigo Image Cytometer (Nexcelom) and flow cytometry. Cells were lifted and dissociated into single cells with Trypsin. Details on the antibodies that were used for flow cytometry are listed in Table 2. The cells were incubated with 0.5 ug /ml final concentration of conjugated antibodies in 1% BSA-PBS for 40 minutes on ice and washing was repeated as before. The cells were then suspended in 300 uL 1% BSA-PBS with DAPI (1:2000) for live/dead separation and kept on ice to be used for the flow cytometry analysis.

**Supplementary Table 2.** Antibodies used for the detection of GFP-OCT4, CD133 and SSEA4 by flow cytometry in the compound screening experiment.

| Antibody | Target/function | Ratio | Number of cells |
| --- | --- | --- | --- |
| CD133-BV786, Mouse Anti-Human, clone W6B3C1, BD 747640 | CD133 | 0.6 μL in 100 μL | 300,000 |
| Mouse IgG1-BV786, BD 563330 | isotype control for CD133 | 0.6 μL in 100 μL | 300,000 |
| Alexa647 Mouse anti-SSEA-4 clone MC813-70 | SSEA4 | 2.5 μL to 100 μL | 300,000 |
| Alexa647 Mouse IgG3 | isotype control for SSEA4 | 0.31 μL to 100 μL | 300,000 |

### Gene knockdown

The stable knockdown of candidates was performed by commercial shRNA constructs (Merck): TRCN0000236473; TRCN0000131081; TRCN0000127634. shRNA plasmid DNA was transfected into cells with Lipofectamine 3000 (Thermo Fischer Scientific) according to manufacturer guidelines. Puromycin was added to the growth media at 0.1ug/ml concentration and individual colonies were picked, expanded, and screened for gene knockdown compared to Scramble control transfected cells. The primers used for qPCR verification for BRD9 were as follows: Forward: CGCAGGCTTTAAGATGATGAGC, reverse: GCTCCTCTGCGGTACTGTC.

### Nucleic acid extraction from cell lines

RNA was extracted using Direct-zol (TM) RNA extraction kit according to manufacturer protocol (Cambridge Bioscience, R2052). The quality of the RNA samples was verified using an RNA screen tape on a Tape-Station (Agilent). The RIN values for all samples were >7.5.

### RNA Isolation and cDNA synthesis

Total RNA was isolated by RNeasy RNA Extraction Kit (Qiagen) according to manufacturer’s guidelines. RNA was then eluted in 30µl of water and the concentration was measured using Nanodrop. The master mix was prepared as follows: 8µl 5x First-Strand Buffer (Invitrogen), 0.5µl Random primers (0.5 µg/ml) (Promega Cat. C1181), 1µl dNTP mix (10 mM each) (Promega Cat.U1515), 2 ul 0.1 M DTT, 0.5 µl RNase Out, 0.25 µl Superscript III Reverse Transcriptase (Life Technologies). 500 ng of total RNA into a separate tube with 11.75 µl RNase-free water. RNA was heated to 65 c for 5 min and allowed to chill on ice for 2 min. 8.25 µl of the master mix were added to RNA. The reaction was incubated at 25 c for 10 min and then at 42c for 50 min. The reaction was then inactivated by heating at 70 for 15 min.

### RT-qPCR

2ng of synthesized cDNA was added to 5µl Power SYBR Mix (Life Tecnologies, 4368708 (Master Mix)) and 1.5µl 2µM of forward and reverse primers. RT-qPCR was performed on ViiA 7 machine with the following intervals: denaturation (95 c) for 15s and a total of 40 cycles, C) for 60s, final extension (60 cC) for 10 minutes.

### Flow cytometry for cell cycle analysis

A13A and FG PDAC cells in which the FUCCI construct was incorporated were taken from adherent conditions and counted, then plated in spheroid conditions at a density of 5,000 cells / 1 mL medium for 10 days, before the cells were collected and analysed using Fortessa (BD Bioscience). Passaging was performed a day 5, after which cells were plated again in spheroid conditions, with the same initial density of 5,000 cells / 1 mL medium. Compounds were added in day 7, and the treatment lasted for 72 hours, with 4 different conditions: I) I-BRD9 (10 μM) ; II) gemcitabine (5 μM); III) I-BRD9 (10 μM) + gemcitabine (5 μM); IV) DMSO control. Experiment was performed in 3 replicates. The data was analysed in FlowJo.

### Preparation and Sequencing of Illumina RNA libraries

RNA-Seq libraries were created using the NEBNext Ultra RNA library prep kit using TruSeq indexes, following the manufacturer’s recommendations. In summary, 500 ng of total RNA was used to isolate mRNA poly(A) by two rounds of purification using oligo dT magnetic beads followed by fragmentation and cDNA synthesis by random primers and reverse transcriptase. Bar-coded adapters were ligated to the cDNA fragments and a PCR reaction was performed to produce the sequencing libraries. To verify the library concentration and the library fragments length was used Agilent 2200 Tape-Station System. Adapter-ligated cDNA fragment libraries were sequenced on a NextSeq 500 platform (Illumina) using a paired-end run (2 × 41 bp).

### RNA-sequencing analysis

Sequencing reads from the RNA-seq experiment were aligned to the human genome (hg38) using HISAT2 with the default parameters. FeatureCounts was used to assign mapped reads to genes with annotation gtf file Ensembl94. Differential gene expression analysis was performed using DESeq2 with IHW method for p value adjustment, apeglm method for effect size shrinkage, FDR-adjusted p value < 0.05 and log2 fold change > 0. Functional analysis of differential expressed genes was performed using Metacore. All plots were generated using R package 3.6.

### Western blot analysis

Protein was isolated by lysing cells with RIPA Buffer (Sigma-Aldrich) supplemented by cOmplete EDTA-free protease inhibitor (Roche) and PhosSTOP^™^ (Sigma-Aldrich) and extracting the supernatant after high-speed centrifugation at 4°C. Protein quantification was performed using the Pierce BCA Protein Assay kit following the manufacturer’s protocol. Isolated proteins were prepared for SDS-PAGE separation by dilution with 4× NuPAGE Sample buffer (Invitrogen), addition of NuPAGE™ Sample Reducing Agent ((10X), Invitrogen), 95°C for 5 minutes, and cooling. Isolated proteins were then analysed by Western blotting. Protein separation via SDS-PAGE was performed on a NuPAGE 4%–12% or 12% Bis-Tris gel (Life Technologies) with NuPAGE™ MOPS SDS Running Buffer (Life Technologies). Proteins were transferred to a PVDF membrane, blocked with 5% milk in PBS and 0.05% tween 20, probed with protein-specific antibodies, incubated with horseradish peroxidase-conjugated secondary antibodies, and visualized via enhanced chemiluminescence using the SuperSignal West Pico Chemiluminescent Substrate (Thermo Scientific). All antibodies (Supplementary Figure 4) were diluted in 5% milk in PBS and 0.05% tween 20. Quantification was performed using ImageJ gel analysis tool.

**Supplementary Table 4.**
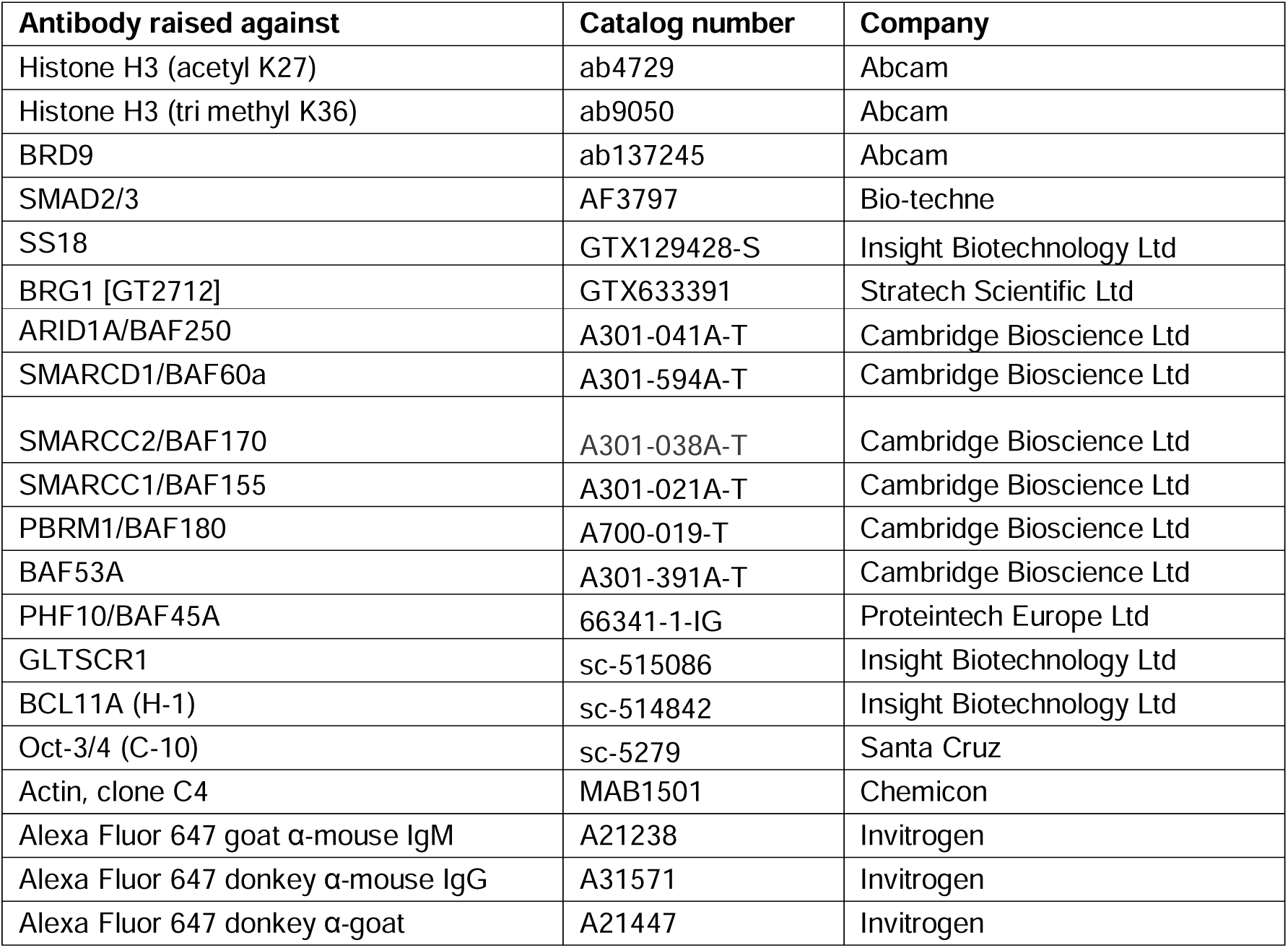
Antibodies used in this study.

### Immunostaining

The immunostaining method has been described previously^3-5^. Cells were fixed for 20 minutes at 4°C in PBS 4% PFA (electron microscopy grade), rinsed three times with PBS, and blocked and permeabilized at the same time for 30 minutes at room temperature using PBS with 10% Donkey Serum (Biorad) and 0.1% Triton X-100 (Sigma). Incubation with primary antibodies diluted in PBS 1% Donkey Serum 0.1% Triton X-100 was performed overnight at 4°C. Samples were washed three times with PBS, and then incubated with AlexaFluor secondary antibodies for 1 hour at room temperature protected from light. Cells were finally washed three times with PBS, and Hoechst (Sigma) was added to the first wash to stain nuclei. Images were acquired using a LSM 700 confocal microscope (Leica).

### Chromatin Immunoprecipitation (ChIP)

All steps were performed on ice or at 4°C and ice-cold buffers and PBS were supplemented with 1mg/ml Leupeptin, 0.2mM PMSF, and 10mM NaButyrate were used unless otherwise stated. Approximately 5×10^6^ cells were used per sample and cross-linked with 1% formaldehyde for 15 minutes. Cross-linking was stopped by incubating samples with glycine at a final concentration of 0.125M for 5 minutes at room temperature, and the cells were washed with PBS followed by pelleting at 250g for 5 minutes. The pellet was re-suspended in 2ml ChIP Cell Lysis Buffer (CLB: 10 mM Tris pH8, 10 mM NaCl, 0.2% NP-40) and incubated for 10 minutes to lyse the plasma membranes. Nuclei were pelleted at 600g for 5 min, lysed in 1.25ml of ChIP Nuclear Lysis Buffer (NLB: 50 mM Tris pH8, 10mM EDTA, 1% SDS) for 10 minutes, and then 0.75ml of ChIP Dilution Buffer (DB: 20 mM Tris pH8, 2mM EDTA, 150mM NaCl, 0.01% SDS, 1% Triton X-100) was added to the samples. Chromatin was sonicated in 15ml Diagenode Bioruptor Pico water bath sonicator with an automated water cooling system, by performing 30 cycles of 30 seconds ON, 45 seconds OFF. This protocol resulted in the homogeneous generation of fragments of 100-400bp. Samples were clarified by centrifugation at 16000g for 10 minutes, and diluted with 3.5ml of DB. After pre-clearing with 10µg of non-immune IgG for 1h and 50µl of Protein G-Agarose for 2h, ChIP was performed overnight in rotation using specific antibodies (Table S2) or non-immune IgG as a control. After incubation for 1 hour with 30µl of Protein G-Agarose, beads were washed twice with ChIP Washing Buffer 1 (WB1: 20mM Tris pH8, 2mM EDTA, 50mM NaCl, 0.1% SDS, 1% Triton X-100), once with ChIP Washing Buffer 2 (WB2: 10mM Tris pH8, 1mM EDTA, 0.25M LiCl, 1% NP-40, 1% Deoxycholic acid), and twice with Tris-EDTA (TE: 10mM Tris pH8, 1mM EDTA). Precipitated DNA was eluted with 150µl of ChIP Elution Buffer (EB: 100mM NaHCO3) twice for 15 minutes at room temperature in rotation, and processed as follows in parallel with 300 µl of sonicated chromatin non-used for ChIP (Input). Cross-linking was reverted by adding NaCl to a final concentration of 300mM for protein-DNA de-crosslinking and incubated at 65°C for 5 hours and 1 µg RNase A (Sigma) to digest contaminating RNA. Finally, 60 µg of Proteinase K (Sigma) were added overnight at 45°C. DNA was extracted by sequential phenol-chloroform and chloroform extractions, and precipitated overnight at -80°C in 100mM NaAcetate, 66% ethanol and 50µg of glycogen (Ambion) as a carrier. After centrifugation at 16,000g for 1 hour at 4°C, DNA pellets were washed once with ice-cold 70% ethanol, and finally air dried. ChIP samples were resuspended in 30µl and 1:10 of the samples were used in qPCR for verifying the ChIP samples.

### Chromatin Immunoprecipitation (ChIP) sequencing

Tagmentation was performed on ChIPed DNA by Tn5 transposase from Nextera DNA Sample Prep Kit (Illumina, cat.#FC-121-1031). Tagmented DNA purified by DNA Clean & Concentrator-5 kit (Zymo research, Cat# 4013). Tagmented DNA was subjected to PCR amplification and double-size selection by AMPure XP beads (Beckman, cat.#A63881) (0.55x/0.9x). The library were multiplexed, quantified using a High-sensitivity d1000 TapeStation (Agilent) and then sequenced using a NextSeq 500 (Illumina) (paired-end, 2 × 41 bp). Sequencing depth was >20 million reads per sample.

### ATAC-sequencing

Cells were washed once with PBS, collected in Cell Dissociation Buffer (Gibco 13150-016) or TrypLE and centrifuged at 300g for 3 min. The cell pellets were then resuspended in 2 ml of 4°C PBS and counted by haemocytometer for using 100,000 cells in the subsequent step. Cells were centrifuged at 300g for 3 min, the supernatant aspirated, the cell pellet resuspended in 150 ul of Isotonic Lysis Buffer (10 mM Tris-HCl pH 7.5, 3 mM CaCl, 2 mM MgCl2, 0.32 M Sucrose and Protease Inhibitors, Roche), and incubated for 12 min on ice. Triton X-100 from a 10% stock was then added at a final concentration of 0.5%, the samples were vortexed briefly and incubated on ice for 6 min. The samples were centrifuged for 5 min at 400g at 4°C, and the cytoplasmic fraction removed from the nuclear pellet. The samples were resuspended gently in 625uL of PBS and transferred to a fresh 1.5 ml eppendorf tube. The nuclei were centrifuged at 1500g for 3 min at 4°C and the supernatant aspirated thoroughly from the nuclear pellet. This step was immediately followed by tagmentation (Nextera DNA Sample Preparation Kit for 24 Samples, FC-121-1030) by resuspending each sample in 100 µL Nextera mastermix (52.5 µl TD buffer, 42.5 µl of water and 5 µl of TDE1 per reaction). The nuclear pellet was resuspended thoroughly by pipetting and incubated at 37 °C for 1 hour shaking at 300rpm. The reaction was stopped with 300 µL of buffer PB from the Qiagen PCR purification kit, followed by Qiagen PCR clean up protocol using MinElute columns and eluting each sample in 18 µl buffer EB. For the control sample, the nuclear pellet was subjected to genomic DNA isolation with GenElute Mammalian Genomic DNA Miniprep Kit (Sigma, G1N70) according to manufacturer’s protocol, and the purified genomic DNA was thereafter immediately used for tagmentation as for other ATAC-seq samples.

Next a PCR reaction (for all samples including control sample) was performed with the following constituents: 10 µl template from tagmentation, 2.5 µl I7 primer (Nextera® Index Kit with 24 Indices for 96 Samples, FC-121-1011), 2.5 µl I5 primer, 10 uL Nudease Free H2O 25µl NEBNext High-Fidelity 2x PCR Master Mix (New England Labs Cat #M054 and 10 uL Nuclease Free H2O. The PCR settings were as follows: at 72 °C for 5 miutes, initial denaturation at 98 °C for 30 seconds, then 12 cycles of 98 °C for 10 seconds, primer annealing at 63 °C for 30 seconds and elongation at 72 °C for 1 minute, and holding at 4 °C. After completing the PCR, the sample were stored at -20 °C. The PCR primers were removed with 1 x 0.9:1 SPRI beads (Beckman Coulter, Cat no. A63880) according to manufacturer’s protocol and samples eluted in 20 µl. 2 µl of the samples were run on Agilent HS Bioanalyzer HS for confirming the size selection of the ATAC libraries. ATAC-sequencing was performed by Illumina HiSeq 2000 sequencing with 75 bp PE for obtaining more than 40 million mapped reads per library. The raw and processed ATAC-seq data are publicly available on GEO (Accession number: GSE222952).

### ATAC-sequencing analysis

Sequencing reads from the ChIP-seq and ATAC seq experiment were aligned to the human genome (hg38) using bowtie with reporting mode,” –best –strata –v2”. Deeptools was used to generate covergae track(bigwig). Coverage track was visualized by using UCSC genome browser. Peak calling was performed by using macs2 peak caller with default parameters for ChIP seq, and with parameter “--nomodel --shift -100 --extsize 200” for ATAC seq. Peaks annotated with nearest gene information by using BEDTools. Peak distribution over different genomic features were summarized by using Bioconductor package ChiPpeakAnno. Motif enrichment analysis within peak regions was performed using HOMER. All plots were generated using R package 3.6.

For the TF footprint analysis, motifs in JASPAR database were scanned using ATAC bam file and peak coordinates for Ctrl and IBRD9 separately. We then compared the cleavage profiles from Ctrl and IBRD9 for each TF: 2027 IBRD9 ATAC peaks within IBRD9 specific loop anchors and 2372 Ctrl ATC peaks within Ctrl specific loop anchors.

### Single-cell RNA-sequencing on patient samples

#### Processing of the patient tumour samples

The Collagenase IV (Stem Cell Technologies, cat no: 7909) aliquot was defrosted in a water bath before sample collection. The freshly surgically removed PDAC sample was kept in the primary growth medium (500ml of DMEM/F12; 1:100 ml of L-glutamine; Pen-strep; 500ul of thermostable FGF2; 20ng/ml of EGF; 5ug/ml of Insulin; 1:50 of B27) after the surgery, and processed further inside a sterile Microbiological Safety Cabinet.

The tissue sample in the primary growth medium was poured onto a sterile 10cm petri dish and the sample was broken up into as small pieces as possible by using a scalpel. By using sterilised forceps, the tissue was collected into the 15ml falcon tube containing 10ml of primary growth medium + 2.5 mg/ml Collagenase IV + 2cmg/ml Dispase II (Sigma-Aldrich, Cat. no. 4942078001) + 1c g/ml Trypsin Inhibitor (Sigma-Aldrich, Cat. no. T6522) + 1 unit/ml DNase I (NEB, Cat. no. M0303S) and incubated at 37°C with shaking speed at 50rpm for about 40 min. The dissociated cells were repeatedly collected at intervals of 20 min to increase cell yield and viability. Cell suspensions were filtered using a 70um cell strainer and red blood cells (RBC) were removed by RBC lysis buffer (Invitrogen, Cat. no. 1966634) with 1 unit/ml DNase I. The digested tissue was decanted into a 15ml falcon with growth medium and spun at 300xg for 10mins to sediment single cells. Cells were resuspended in primary growth media, counted (live/dead) and divided into four treatment conditions: i) DMSO. ii) I-BRD9 (Bio-Techne (R&D Systems), Cat no. 5591/10) at 10µM final concentration. iii) DMSO + Gemcitabine at 1µM final concentration. iv) I-BRD9 + Gemcitabine. Cells were transferred to low-adherent flat-bottomed plates. Cells were incubated with four different conditions for 72 hours. Cells were collected for single-cell RNA-sequencing.

#### Preparation of Single Cell Samples

After collection, the cells were centrifuged at a speed not exceeding 400rcf. The supernatant was discarded. The cell pellet was resuspended in 1mL 1X PBS containing 0.04% BSA and the washing procedure was repeated twice. After washing, appropriate volume PBS was added to the cell precipitation to obtain single-cell dispersion suspension with a concentration close to the goal number. A wide-bore pipette tip was used for pipetting cell resuspension for lower cell damage. 40µm Cell Strainers were used for removing cell debris and cell clumps. Automatic cytometry was used to determine the cell concentration. The sample volume was calculated based on the optimal cell sampling concentration supplied by the 10X official website and the target capture number. If the calculated concentration was too high, the liquid volume was adjusted to the appropriate concentration and the counting was repeated. Once the desired cell suspension was obtained, it was immediately placed on ice for subsequent GEMs preparation and reverse transcription.

#### Library Construction

The library construction was performed as follows: (1) Interrupt, end repair and add A base. Finally, the SPRI select beads were used to purify the product. (2) Adaptor ligation and SPRI select purification. (3) Index PCR and SPRI select purification. (4) Qubit® 3.0 Fluorometer (Life Technologies) was used to determine the library concentration. (5) Agilent 2100 High Sensitivity DNA Assay Kit (Agilent Technologies) was used to determine the distribution of library product fragments. After the library construction, Agilent 2100/LabChip GX Touch was used to detect the fragment length distribution of the library. Also, qPCR was used to accurately quantify the effective concentration of the library. The effective concentration of the library aimed > 10 nmol/L.

#### Single-cell gene expression analysis

Sequencing data was aligned and quantitated using CellRanger (10x Genomics) and hg38 reference genome sequence. Raw gene expression matrices were then analysed using Seurat R package (v4.0.5). Quality control was performed individually for each sample. All cells with <500 detected genes were removed, as well as cells contained <600 unique molecular identifiers (UMIs) and >10% mitochondrial counts. Raw read counts were normalized using *NormalizeData* function, based on which 2000 variable genes were identified. 2000 genes across samples for integration were selected using SelectIntegrationFeatures function. To perform data integration using reciprocal PCA method in FindIntegrationAnchors function, data scaling and PCA were performed on each sample first. 50 dimensions were used in integration process.

In order to identify cell types, we performed principal component analysis (PCA) on integrated data with default parameters in Seurat to reduce dimensionality. The first 50 principal components (PCs) were used for Uniform Manifold Approximation and Projection (UMAP) dimensionality reduction. The *FindNeighbours* (30 nearest neighbours) and *FindCluster* function (resolution = 2.5) were used to cluster cells. Clusters in 2D UMAP were used to identify cell types based on marker genes. Unless specified, default parameters were used for each function.

#### ChIP-sequencing data analysis

The adapters were removed and the filtered reads were mapped to hg38 reference genome using bwa^6^ mem. The duplicated reads were removed using Picard (https://broadinstitute.github.io/picard/) and the reads with quality lower than 20 were filtered. Bam files were converted to bigwig format file using deepTools^7^ bamCoverage, The parameter ‘scaleFactor’ for each sample were determined by edgeR^8^ function calcNormFactors using “TMM” normalization method. The ChIP-seq peaks were called using the default (narrow) setting in MACS2. The promoter regions were defined as +/-2.5Kb around the TSSs regions.

### *In-situ* ChIA-PET and data analysis

Cells were grown to 80% confluency, and treated by 1% formaldehyde for 20 min and ethylene glycol bis (EGS) (Thermo Fisher Scientific, cat.#21565) for 45 min. 10 million of dual crosslinked A13A CSC cells were used to generate one in-situ ChIA-PET library. Briefly, after cell lysis and nuclear lysis, cells were subjected to in-situ digestion with Alu I restriction enzyme for overnight. On the next day, A-tailing was performed using Klenow Fragment (3′>5′ exo-) (NEB, cat.#M0212L) and dATP (100 mM; NEB, cat.#N0440S) for 1 hr at room temperature. In-situ ligation was performed using in-house bridge linker (F: 5′-/5Phos/CGCGATATC/iBIOdT/TATCTGACT -3′, R: 5′-/5Phos/GTCAGATAAGATATCGCGT -3′. HPLC purified, from Integrated DNA Technologies) and T4 DNA ligase (Thermo Fisher Scientific, cat.#EL0013) for RT 1 hr and then 16°C for overnight. The nucleus was sonicated into ∼1 kb size. Then chromatin was precleared using protein G beads for 1 hr and immunoprecipitation was performed using anti-H3K27ac antibody (Active Motif, cat.# 39133) coated protein G beads (Life Technologies, cat. no. 10009D) for overnight.

After that, proximity-ligated chromatin complexes were washed sequentially by low salt, high salt, LiCl buffer and TE buffer. Chromatin complexes were eluted from the beads by incubation with ChIP elution buffer (1% SDS+TE) at 65 °C for 1 hr. After that, proteinase K (Life Technologies, cat.#AM2548) was added to the elution reverse cross-linked chromatin complexes for overnight. After DNA purification using QIAquick PCR Purification Kit, tagmentation was performed on the proximity-ligated DNA by Tn5 transposase from Nextera DNA Sample Prep Kit (Illumina, cat.#FC-121-1031). Then tagmented DNA was immobilized on M280 streptavidin dynabeads (Invitrogen, cat.#11205D). PCR amplification was performed on beads using Nextera DNA Sample Prep Kit and the products were purified by AMPure XP beads (Beckman, cat.#A63881) and subjected to size selection (350-600 bp) on BluePippin instrument (Sage Science) using Blue Pippin Cassette Kit (Sage Science, cat.#BDF2010). DNA library was sequenced on Illumina Novaseq 6000 by paired-end 150 bp.

### *In-situ* ChIA-PET data processing

We used ChIA-PIPE^9^ to process ChIA-PET data. Pair-end reads containing bridge linker were kept and trimmed, the flanking sequences were mapped to human hg38 reference genome and uniquely aligned pair-end tags (PETs) with MAPQ ≥ 30 were retained. Redundant reads were then removed and bam format files were used for downstream analysis. Each PET was classified as self-ligation PET or inter-ligation PET using a genomic span cutoff of 8Kb. Inter-ligation PETs were extended by 500 bp on both ends and clustered. The PET clusters with counts lower than 10 and spanning less than 5 Kb or larger than 1Mb were filtered. We referred the remaining PET clusters as chromatin interaction loops. Un-clustered individual inter-ligation PETs were referred as PET singletons. All PETs (including self-ligation and inter-ligation PETs) were used to call peaks using MACS2^10^.

Chromatin interaction loops from all samples were merged to form a union set. Read counts for each merged loop were counted across all samples to form a loop – sample count matrix. To identify control-specific and IBRD9-specific loops, the read matrix was imported into R and a negative binomial model was fitted using glmFit function in edgeR^8^ R package. The loops with p-value <0.1 were determined as differential loops^11^. Loop anchors proximal to (less than 3Kb) known transcription start sites (TSSs) were defined as promoters and the remaining loop anchors were defined as distal regulatory elements.

### Functional enrichment of downregulated genes with IBRD9 treatment

Functional enrichment of genes network was generated using the ToppFun application of the Toppgene Suite using a corrected (Benjamini and Hochberg) p-value cut-off of 0.05 (https://toppgene.cchmc.org/)^12^. Cytoscape application (https://cytoscape.org/)^13^ was used to generate the functional enrichment network.

### Visualization

Statistical plots were created using R. Screenshots of chromatin interactions was visualized using WashU Genome Browser.

### Animal studies

All research protocols conformed to the 2011 Guidelines for the Care and Use of Laboratory Animals published by the National Institutes of Health. All animal use protocols and experiments were performed according to the guidelines approved by the Institutional Animal Care and Use Committee at Guangdong Medical Laboratory Animal Center and Yat-sen University. All NSG mice (Model Animal Research Center of Nanjing University) and PDX models (PDXM-221Pa, Shanghai Medicilon Inc) were maintained in a specific pathogen-free facility with high efficiency particulate air (HEPA)-filtered air and given autoclaved food and water.

### Establishment of xenograft tumor models in nude mice

The A13A cells for subcutaneous injection were divided into the following groups, 1) Null group, cells were transduced with scrambled shRNA adenovirus (Ad-GFP-U6-shScram, Vector Biolabs) containing a scramble sequence under the control of U6 promoter, with the GFP co-expression under a cytomegalovirus (CMV); 2) BRD9 KD group, cells were transduced with BRD9-targeted shRNA adenovirus (Ad-GFP-U6-shBRD9, Vector Biolabs); 3) Gemcitabine group, cells were treated with gemcitabine (7.5 μM, Selleckchem, LY-188011); 4) BRD9 KD + Gemcitabine group, cells transduced with Ad-GFP-U6-shBRD9 were treated with gemcitabine additionally. These cells were maintained at a low-density culture (1250 cells/ml, 312.5 cells/cm2) and harvested for subsequent experiments 72 h post transduction/treatment.

Tumor studies were performed in female 8-10 weeks old NSG mice. For tumor formation, 0.5×10^6^ cells in 200µl of Matrigel/DMEM media mixture (1:1 ratio) were subcutaneously injected into the right flank of NSG mice.

### Tumor growth and mouse survival

For tumor growth, mice were monitored twice a week or every 3 days by observation and palpation. Tumor size was measured with a digital caliper and tumor volume was calculated using the formular LxW2×0.5. Mouse body weight was measured twice weekly. Death, weight loss of 15% or more of body wight, tumor volume with 2,000m^3^ or more, or severe labored breathing or dyspnea were considered endpoints and all mice were euthanized.

### In vivo PET imaging

Mice were fasted for 12 h before 18F-fluorodeoxyglucose (FDG) injection but allowed free access to water. After anaesthesia with 2% isoflurane, mice were injected with 18F-FDG (11 to 16MBq, German Cancer Research Center, Heidelberg, Germany) by tail vein and kept at 37oC until imaging. Imaging was started at 60 minutes after 18F-FDG injection with non-invasive micro-positron emission tomography (μPET, Siemens Inveon). A CT scan (80 kVp, 500cμA, at 120 projections; approximately 4-minutes) was acquired for anatomical reference and enabled PET attenuation correction during reconstruction. Postprocessing was carried out with Inveon Research Workplace.

### Therapeutic studies in PDX models

Therapy was initiated in PDX mice after the tumor reached an average volume of 150mm^3^ with a repeated injection schedule (once every 3 days for a total of six treatments within 15 days). The PDX mice with various treatment were divided into the following groups, 1) Control treatment group (Ctrl), the mice were administrated with DMSO in the same manner as the treatment groups; 2) Gemcitabine treatment group (Gem), the mice were administrated with Gemcitabine (50 mg/kg i.p.); 3) Combination of BRD9 inhibitor with Gemcitabine (IBRD9 + Gem), the mice were administrated with Gemcitabine (25 mg/kg i.p.) and BRD9 inhibitor (10 mg/kg i.p.).

### Histology analysis with H&E staining

Mouse tumor tissues were dissected, rinsed with PBS and fixed in 10% formalin for 48 h. Following dehydration through a series of ethanol solutions, the tissues were embedded in paraffin wax according to standard laboratory procedures. Subsequently, they were sectioned (5 µm thick) using a microtome. Following deparaffinization and rehydration, H&E staining was performed on the sections. In brief, the sections were stained in Mayers Hematoxylin for 1 min. Following rinsing in tape water, the sections were stained in Alcoholic-Eosin for 1 minute and dehydrated and cleared with xylene. The images were examined with an Olympus BX41 microscope equipped with a CCD (Magna-Fire TM) camera.

### Immunohistochemistry assay

After antigen retrieval in sodium citrate buffer (10mM Sodium Citrate, 0.05% Tween 20, pH 6.0), the tumor frozen sections were incubated for overnight at 4°C with primary antibody (anti-Ki67, Abcam, ab15580, 1:200) in 0.02% Triton. After triple washing in PBS, slides were incubated for 45 min at 37 °C with fluorescence conjugated second antibodies (Jackson Immuno Research). DAPI was used for nuclear counterstaining. Four fields of each section were examined for quantification. Fluorescent imaging was performed with an Olympus BX41 microscope equipped with an epifluorescence attachment.

### ChIP-qPCR of anchor regions

The cells were treated as for RNA-sequencing experiment and the ChIP was performed as described above. The primers used for ChIP-qPCR of anchor regions are listed in Supplementary Table 5.

**Supplementary Table 5.**
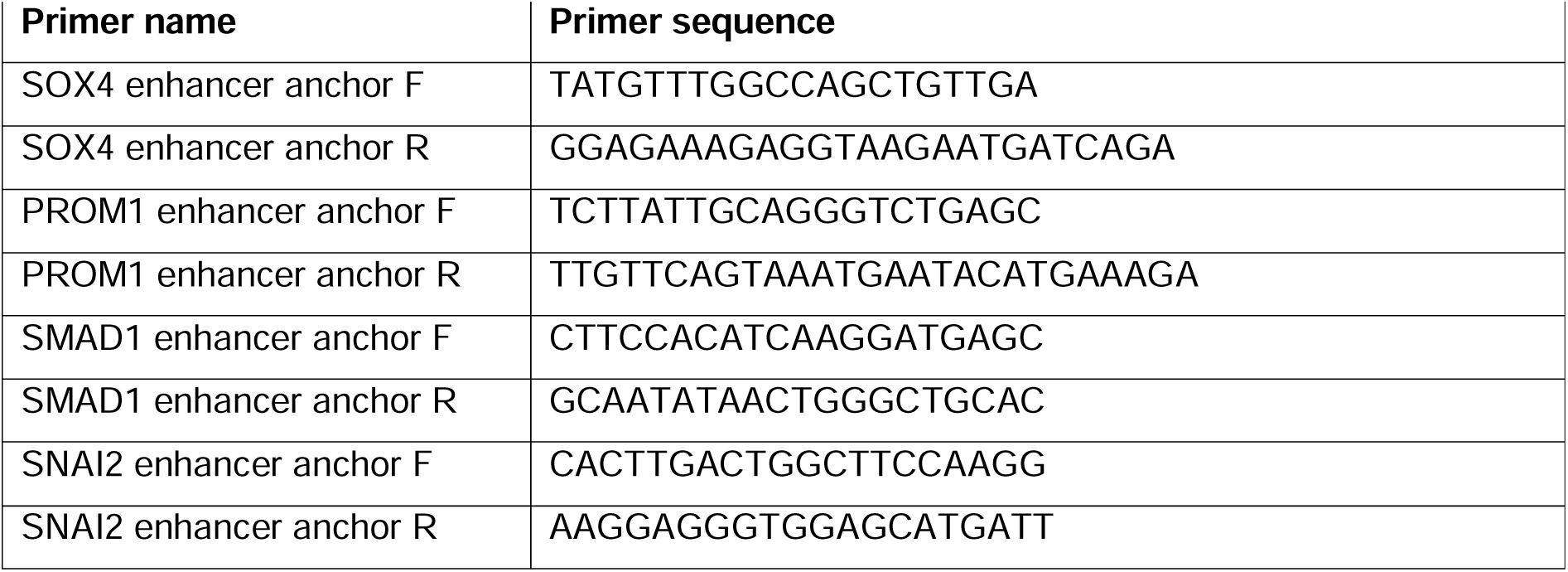
ChIP-qPCR primers of anchor regions.

## Supplemental Figures

**Supplemental Figure 1. BRD9 inhibition by small molecules or knockdown reduces CSCs.** (**A**) Establishing the cell line platform for small molecule screening. eGFP ORF was inserted in frame at the 3’ region of the endogenous OCT4 locus via TALEN-mediated recombination and selected for the insertion with Puromycin that is inserted with eGFP under the control of the PGK promoter. The insertion of eGFP into the OCT4 locus of FG cells was validated by PCR amplification of 5’ and 3’ regions by using primers covering the OCT4 locus and the eGFP sequence in the correct orientation. (**B**) TGFβ/Activin signalling induces chemoresistant OCT4-GFP+/CD133+/SSEA4+ CSCs. Cells expressing CSC markers OCT4-GFP, CD133 and SSEA4 are more resistant to Gemcitabine, 5-FU and Paclitaxel treatment than CSC marker negative cancer cells. FG cells were treated with 0.5µM Gemcitabine, 3µM 5-FU and 0.5µM Paclitaxel in combination with Activin A or 10 µM SB431542 for 5 days, followed by flow cytometry analysis of CSC markers OCT4-GFP, CD133 and SSEA4. Cells were gated for DAPI negative signal for live cells. While most cancer cells are killed by the chemotherapy reagents, the DAPI negative surviving cells are enriched for cancer stem cell markers. (**C**) TGFβ/Activin-SMAD2/3 signalling improves cell survival upon the treatment with chemotherapy reagents. Cells were treated with Gemcitabine, 5-FU and Paclitaxel as in (**B**). (**D-E**) Higher expression of CD133/PROM1 (p-value=0.0292) and OCT4 (p-value=0.135) correlates with lower survival of pancreatic cancer patients according to TCGA data (**D**), and lower disease-free survival (**E**). (**F**) Schematic depiction of BRD9 PROTAC that leads to the degradation of BRD9 protein. (**G**) dBRD9 PROTAC mediated BRD9 protein degradation. Western blotting of BRD9 indicates the reduction of BRD9 protein level upon 10µM or 50µM dBRD9 treatment for 24h or 72h. (**H-I**) BRD9 stable knockdown clone validation at the mRNA level in FG cells (H) and protein level by western blotting in FG cells (I). (**J-K**) BRD9 stable knockdown clone validation at the mRNA level in A13A cells (J) and protein level (K) by western blotting in A13A cells. (**L**) TGFβ/Activin signalling inhibition by 10 µM SB431542 for 5 days eliminates OCT4-GFP+/CD133+/SSEA4+ CSCs. FG cells were treated with 10 µM SB431542 for 5 days, followed by flow cytometry analysis of CSC markers OCT4-GFP, CD133 and SSEA4. (**M**) The percentage of CSC marker positive cells is reduced by I-BRD9 or SB431542. (**N**) TGF β/Activin signalling inhibition and BRD9 inhibition impair CSC self-renewal and sensitize CSCs for Gemcitabine-mediated destruction in FG cells. (**O**) BRD9 inhibition sensitizes PDAC cells for elimination by Gemcitabine. A13A cells were treated for 5 days and quantified for cell numbers. (**P-R**) BRD9 knockdown abolishes sphere formation by sensitizing them for Gemcitabine and 5-FU treatment in FG cells (**P**) and A13A (**R**). Statistical analysis was performed by 2-way ANOVA with multiple comparisons with Tukey correction and **** marks adjusted P-value <0.0001, *** is adjusted P-value <0.001, ** is adjusted P-value <0.01, * is adjusted P-value <0.05.

**Supplemental Figure 2. BRD9 inhibition impacts G0 phase entry of CSCs and forms a protein complex with SMAD2/3.** (**A-B**) Cell cycle analysis with the FUCCI system in (A) A13A and (B) FG cells, treated with I-BRD9, Gemcitabine or the dual treatment with Gemcitabine and I-BRD9 for 5 days. (**C-D**) Flow cytometry analysis of the FUCCI signal and density blots of (C) A13A and (D) FG cells. (**E**) The protein interaction network based on STRING database analysis depicting SMAD2/3 interaction with the subunits of the BAF protein complex. (**F**) Schematic depiction of the different subunit composition of non-canonical BAF, esBAF and npBAF. SMAD2/3 proteins interact with the subunits of non-canonical BAF, esBAF and npBAF complexes in pancreatic CSCs. Co-immunoprecipitation of unique subunits that are specific for esBAF (BCL11a), npBAF (BAF180) and ncBAF (GLTSCR1).

**Supplemental Figure 3. Inhibition of BRD9 abolishes PDAC tumour growth in mice.** (**A**) Schematic depiction of the mouse experiments. (**B**) Tumour volume measurements indicate reduced tumour sized upon BRD9 knockdown. (**C**) PET imaging of tumours in mice indicates reduced tumour sizes upon BRD9 knockdown and absence of visible tumours upon BRD9 knockdown with Gemcitabine treatment. The images show PET scan in 2D and in 3D with the tumour highlighted in yellow dotted areas. (**D**) Tumour size comparison in different treatments. Tumours from non-treated cells are larger compared to Gemcitabine treated and BRD9 knockdown PDAC cells, whereas BRD9 knockdown with Gemcitabine treatment resulted in undetectable tumours at 3 months of growth. (**E**) Relative tumour weight measurements in the different treatment conditions. (**F**) Histological analysis of tumours. N=necrotic areas.

**Supplemental Figure 4. BRD9 inhibition eliminates CSCs in patient-derived primary PDAC tumor cells.** (**A**) Enriched GO terms for genes that are downregulated in I-BRD9 treated and I-BRD9/Gemcitabine treated sample compared to DMSO and Gemcitabine treated samples. (**B**) Enriched Pathway terms for genes that are downregulated in I-BRD9 treated and I-BRD9/Gemcitabine treated sample compared to DMSO and Gemcitabine treated samples. (**C**) BRD9 inhibition leads to the elimination of cells that are enriched in CSC population such as JUN, SOX4 and SKIL. Expression levels of JUN, SOX4 and SKIL in the CSC population is shown as dot plot graphs where blue colour indicates higher expression level compared to gray colour. (**D**) Heatmap of the lost and gained CREs upon BRD9 inhibition.

**Supplemental Figure 5. BRD9 inhibition changes the enhancer-promoter connectome of CSCs.** (**A**) GSEA analysis of genes regulated by BRD9 inhibition indicate the enrichment of genes involved in epithelial to mesenchymal transition, cell migration, regulation of cell population proliferation and vasculature development. (**B**) BRD9 inhibition leads to the loss of enhancer-promoter looping and loss of expression at stem cell and EMT loci such as CD133/PROM1, SMAD1 and SNAI2. The genomic loci show H3K27ac abundance and gene transcription together with 3D chromatin interactions. (**C-D**) Transcription factor footprint analysis identified SMAD3 footprints based on JASPAR motif database at the ATAC-seq peaks of the anchor regions. (**C**) MA0513 has 254 hits in Ctrl-specific ATAC peaks and 211 hits in I-BRD9-specific ATAC peaks on anchors. (**D**) MA1622 has a total of 1370 hits in Ctrl-specific ATAC and I-BRD9-specific ATAC peaks on anchors. The depth of the footprinting is not statistically different in control and BRD9-I treated samples. (**E**) TGF β/Activin signalling leads to the loss of SMAD2/3 and BRD9 binding to the 3D chromatin looping anchors at enhancers of stem cell and EMT loci in CSCs. ChIP-qPCR of distal anchor regions at SOX4, CD133/PROM1, SNAI2, and SMAD1 loci in CSCs +/-SB431542. (**F**) Schematic depiction of the function of SMAD2/3 and BRD9 in regulating the chromatin looping between enhancer and promoter regions near stemness genes, EMT and invasiveness genes.

