## Supplementary figures for "BRD9-SMAD2/3 orchestrates stemness and tumorigenesis in pancreatic ductal adenocarcinoma"

### Supplemental Figure 1

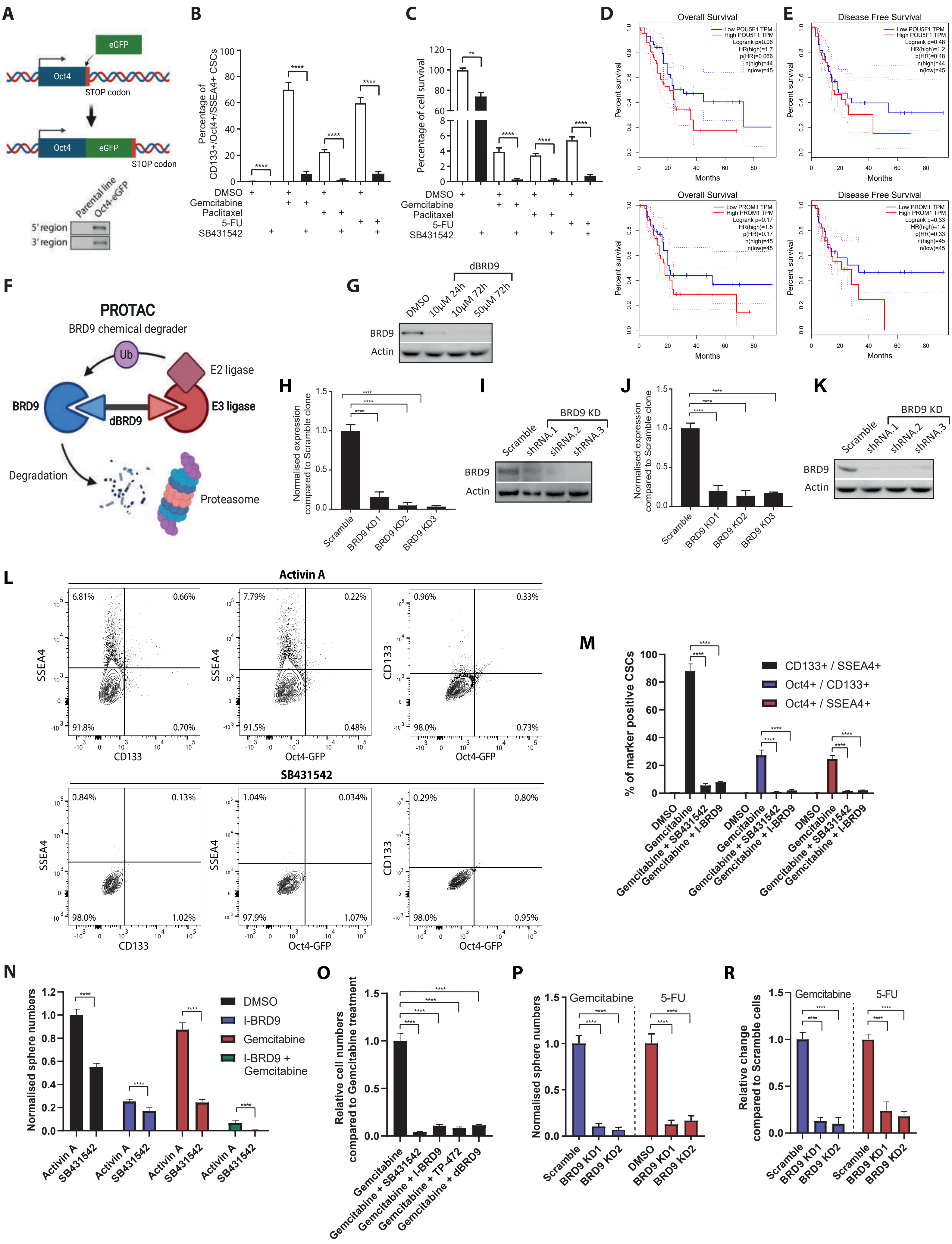

### Supplemental Figure 2

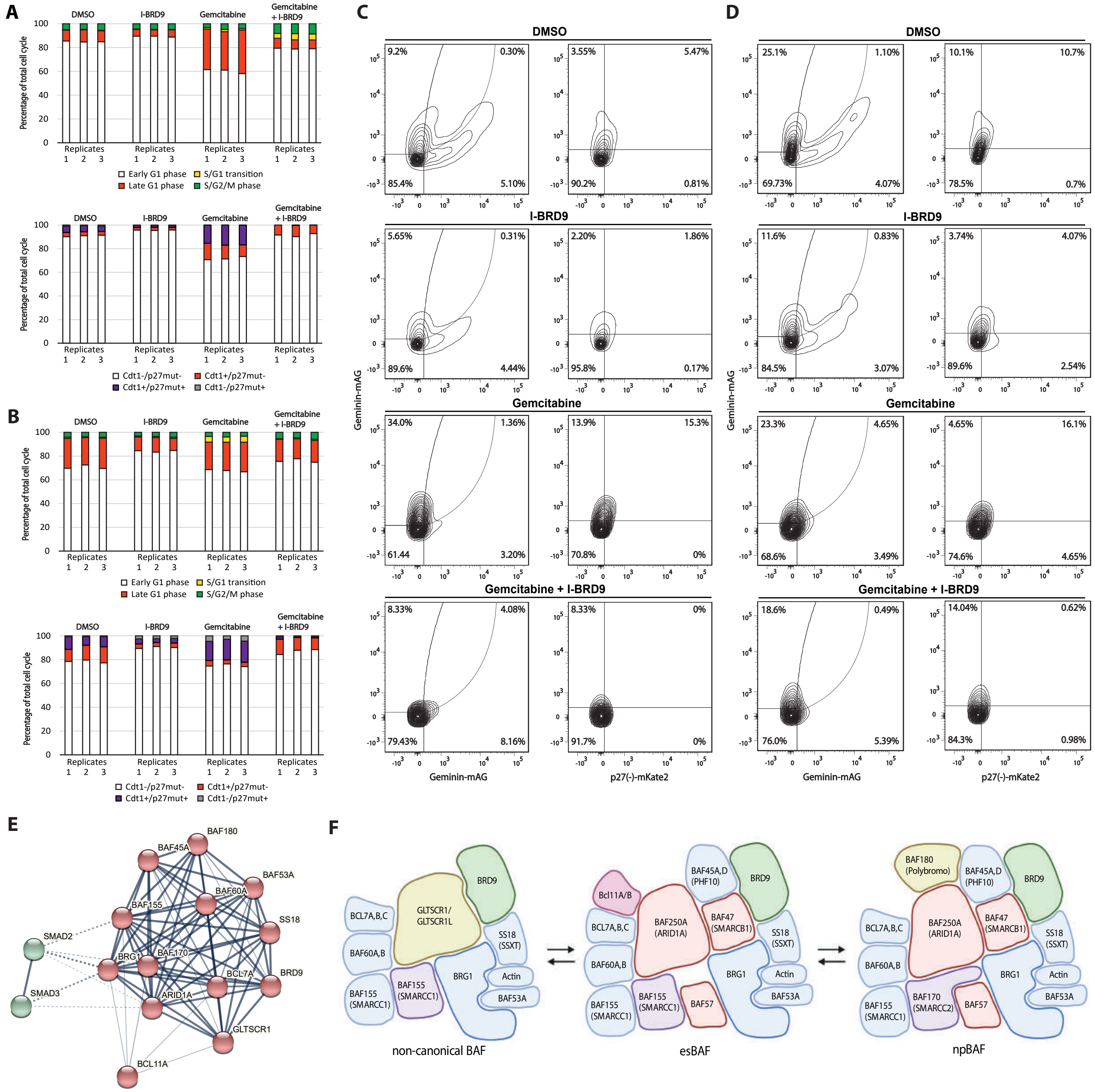

Supplemental Figure 3

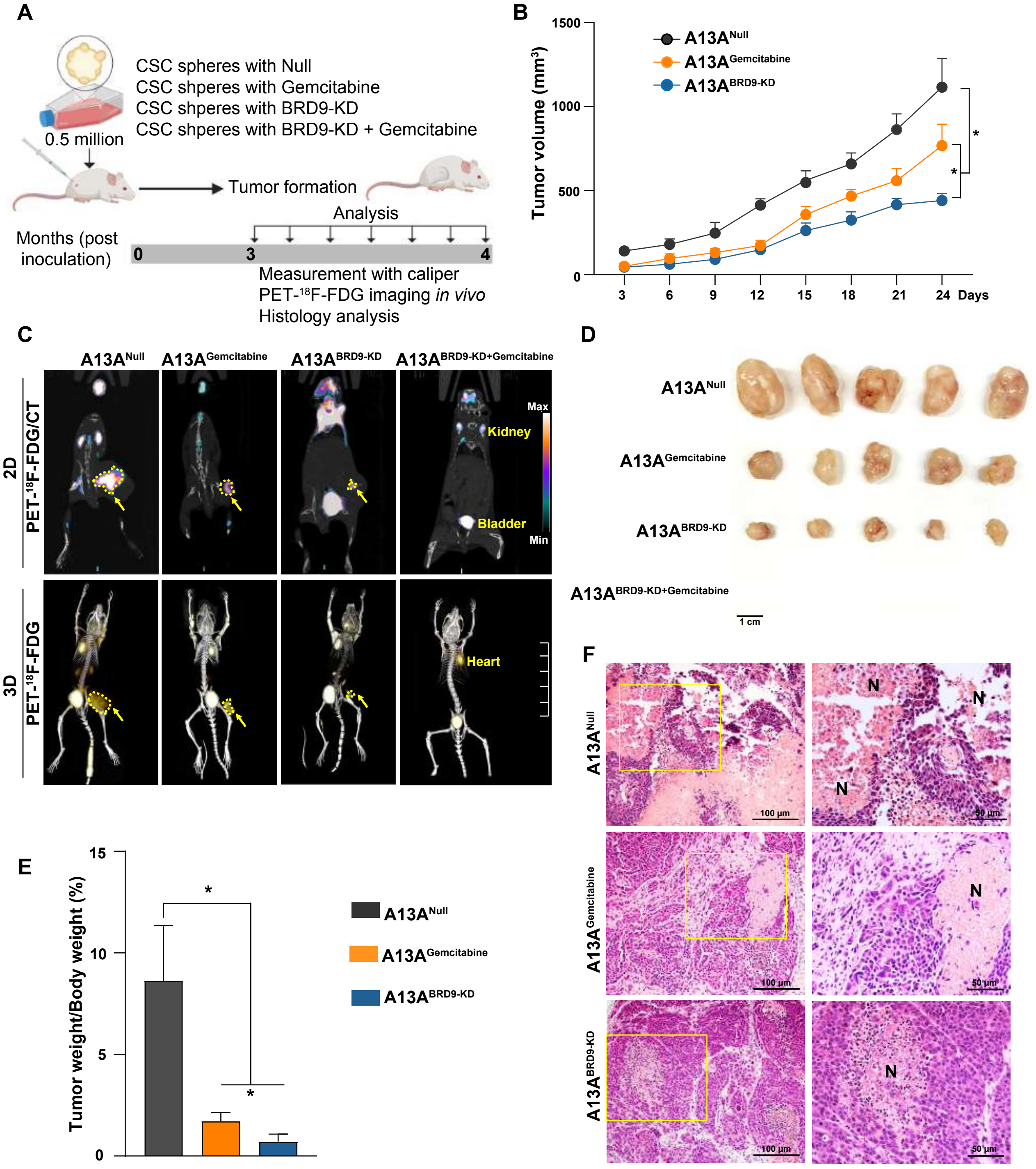

### Supplemental Figure 4

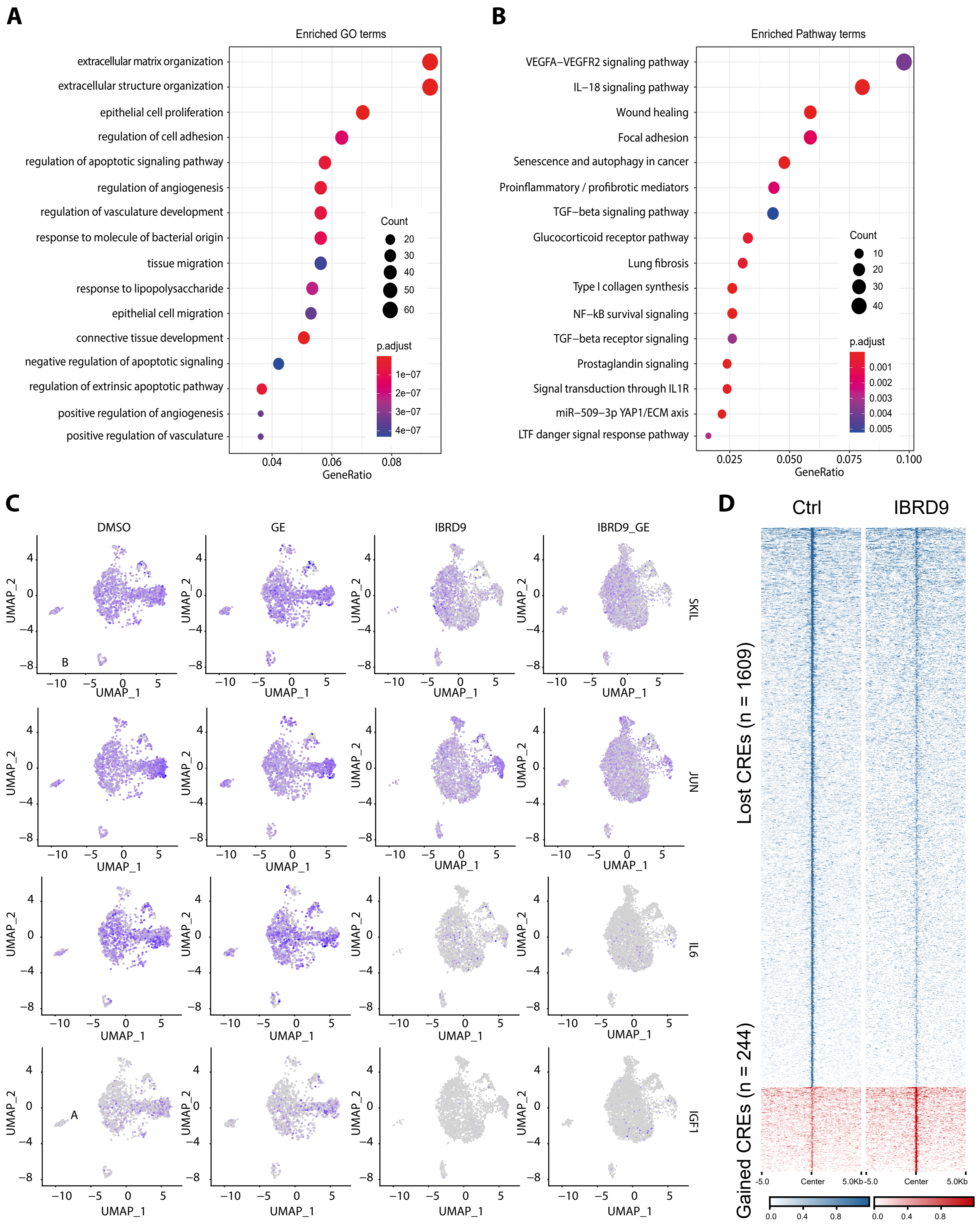

### Supplemental Figure 5

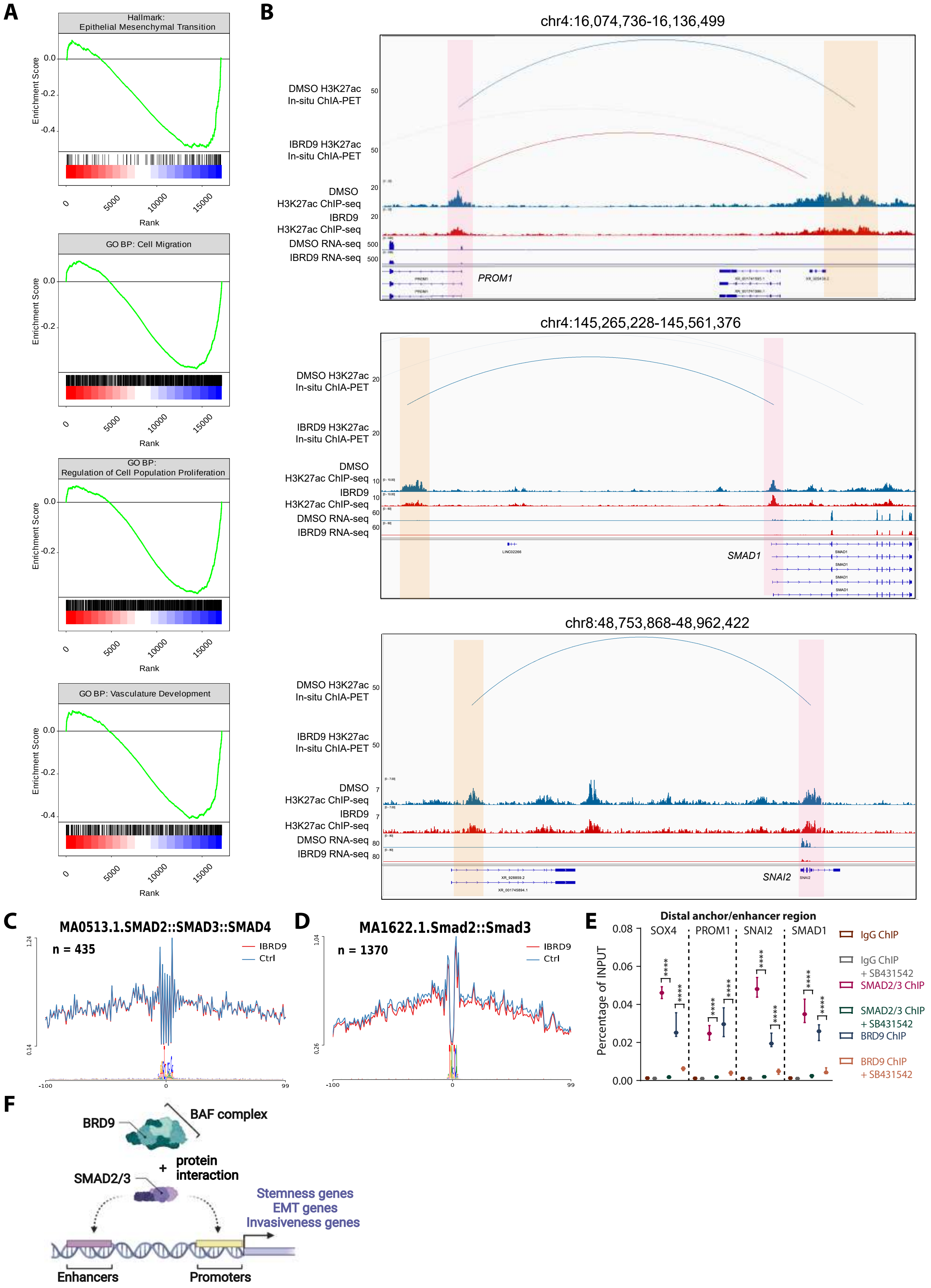
